# A transient dermal niche and dual epidermal programs underlie sweat gland development

**DOI:** 10.1101/2023.04.15.537037

**Authors:** Heather L. Dingwall, Reiko R. Tomizawa, Adam Aharoni, Peng Hu, Qi Qiu, Blerina Kokalari, Serenity M. Martinez, Joan C. Donahue, Daniel Aldea, Meryl Mendoza, Ian A. Glass, Birth Defects Research Laboratory (BDRL), Hao Wu, Yana G. Kamberov

## Abstract

Eccrine glands are mammalian skin appendages indispensable for human thermoregulation. Like all skin-derived appendages, eccrine glands form from multipotent progenitors in the basal skin epidermis. It remains unclear how epidermal progenitors progressively specialize to specifically form eccrine glands, precluding efforts to regenerate these vital organs. Herein, we applied single nucleus transcriptomics to compare the expression content of wildtype, eccrine-forming mouse skin to that of mice harboring a skin-specific disruption of *Engrailed 1 (En1)*, a transcription factor that promotes the formation of eccrine glands in both humans and mice. We identify two concurrent epidermal transcriptomes in the earliest eccrine anlagen: a predominant transcriptome that is shared with hair follicles, and a vastly underrepresented transcriptome that is *En1*-dependent and eccrine-specific. We demonstrate that differentiation of the eccrine anlage requires the induction of a transient and transcriptionally unique dermal niche that forms around each developing gland in humans and mice. Our study defines the transcriptional determinants underlying eccrine identity in the epidermis and uncovers the dermal niche required for eccrine developmental progression. By identifying these defining components of the eccrine developmental program, our findings set the stage for directed efforts to regenerate eccrine glands for comprehensive skin repair.

## Introduction

Eccrine sweat glands are the most numerous appendage found in human skin and are critically important for human thermoregulation, removing excess heat and protecting our bodies from lethal overheating^1–3^. Despite their importance, there are no therapeutic paradigms to regenerate eccrine glands for reconstructive skin repair, leaving patients with epithelial injuries such as extensive burns with extreme, even life-threatening deficits in thermoregulation^1, 2, 4–7^. Thwarting these efforts is a lack of information on how the skin forms these critical appendages in the first place.

Eccrine glands belong to the ectodermal appendage organ class, which also includes hair follicles and mammary glands^2, 8^. These organs form from thickenings of the deepest (basal) layer of the epidermis called placodes, and mature by growing down and differentiating into the underlying connective tissue dermis^8–11^. To date, studies investigating eccrine gland development have largely implicated molecular regulators and developmental pathways shown to also be required for the formation of other ectodermal appendages, like hair follicles and mammary glands^5, 8, 10, 12–18^. However, the molecular factors conferring eccrine identity to epidermal progenitors as they transition through development are poorly understood^5, 8, 10, 12, 19, 20^.

For other major ectodermal appendages such as hair follicles, teeth, and mammary glands, a major function of the placode is to induce local condensation of the underlying dermis to form a specialized niche that engages in coordinated and reciprocal interaction with the appendage progenitors to direct further development^3, 33–37^. Strikingly, eccrine gland placodes, and indeed developing eccrine glands at any stage, are not associated with the morphologically-evident mesenchymal condensations that are the hallmarks of known dermal niches and an analogous niche for the eccrine gland has not been identified^23, 31^. Accordingly, it is unknown if eccrine developmental progression requires extrinsic input from the dermis and adheres to or diverges from the classical ectodermal appendage paradigm^8, 10, 12, 21–25^.

To enable precise targeting on the eccrine gland developmental program, we took advantage of an *in vivo* system that allows direct comparison within the same spatiotemporal context between the developmental program for eccrine glands to that of another major ectodermal appendage, the hair follicle. Specifically, we capitalized on the fact that in the eccrine-forming skin of mice, the palmar/plantar (volar) paw, expression of the *Engrailed 1* (*En1*) gene promotes the formation of eccrine glands and concomitantly inhibits the formation of hair follicles^5, 12, 19, 26–28^. Using genetic modulation of *En1* levels in this tissue, we shifted the composition of mouse volar appendages from eccrine to hair-forming and applied single nucleus transcriptomics to parse out the transcriptional and cellular determinants that make the developing eccrine gland distinct.

## Results

### Targeted disruption of eccrine development by inhibition of eccrine placode identity

*En1* is expressed throughout the deepest (basal) layer of the epidermis in the palmar/plantar (volar) skin of the mouse paw and is focally upregulated in the earliest eccrine anlagen, or placodes, that form therein^12, 19, 29^. This expression pattern is recapitulated in human fetal skin in regions where eccrine glands are developing^12, 28^. Importantly, the level of epidermal *En1* during the developmental period when eccrine placodes are specified is directly correlated with the number of eccrine glands that form in mouse and human skin, and reducing *En1* levels results in a dosage-dependent reduction in the number of eccrine glands^5, 12, 19, 27, 28, 30^. We therefore reasoned that disrupting epidermal *En1* expression at the eccrine placode stage would lead to a specific depletion of the eccrine developmental signature from the skin.

To this end, we derived doxycycline inducible, epidermis-specific, *En1* null, or knock-out, mice (En1-cKO) by breeding mice harboring alleles for a tetracycline responsive Cre recombinase (tetO-Cre); a basal keratinocyte-specific reverse tetracycline transactivator (Krt5-rtTA); and a conditional *En1* null allele (*En1^flox^*)^31–33^. We induced *En1* disruption in the Keratin 5 (Krt5)-positive, basal epidermis by administering doxycycline to pregnant dams on embryonic day (E) 16.5 (Figure 1A). At this developmental stage the first eccrine placodes are forming in the six thickened elevations, or footpads, located at the periphery of the volar hindpaw ^10–12, 19^. On postnatal day (P) 2.5, we harvested the volar skin from En1-cKO (tetO-Cre/+;Krt5-rtTA/+;En1^flox/flox^) and Control animals (En1^flox/flox^, Krt5-rtTA/+;En1^flox/flox^, and tetO-Cre/+;En1^flox/flox^) and compared their volar appendage compositions. Consistent with our previous findings in wildtype mice, developing eccrine glands at multiple stages are simultaneously present in P2.5 Control volar skin^19^. In the footpads, where eccrine gland development began days earlier, nascent-stage eccrine glands that are co-positive for the pan-appendage marker EDAR and for *En1* expression, are differentiating and proliferating into the underlying dermis (Figure 1B)^11, 19^. Importantly, the centrally-located interfootpad volar skin of Control mice is populated by placode-stage eccrine glands that show characteristic upregulation of *En1* and are also positive for EDAR (Figure 1B-bottom left)^10, 11, 19^. In contrast, the footpads and the interfootpad regions of P2.5 En1-cKO mice are significantly depleted of nascent eccrine glands and eccrine placodes, respectively (Figure 1B right panels). Moreover, we find that the interfootpad regions of En1-cKO mice are populated by hair follicle placodes (Figure 1B, bottom-right). Each hair follicle placode is readily differentiated from its eccrine counterparts by an underlying dermal condensate (DC), a morphologically evident aggregate of dermal cells that is the instructive niche for each developing hair follicle and the precursor of the dermal papilla (Figure 1B, bottom-right, arrowheads indicate DC)^8, 21^.

**Figure 1.**
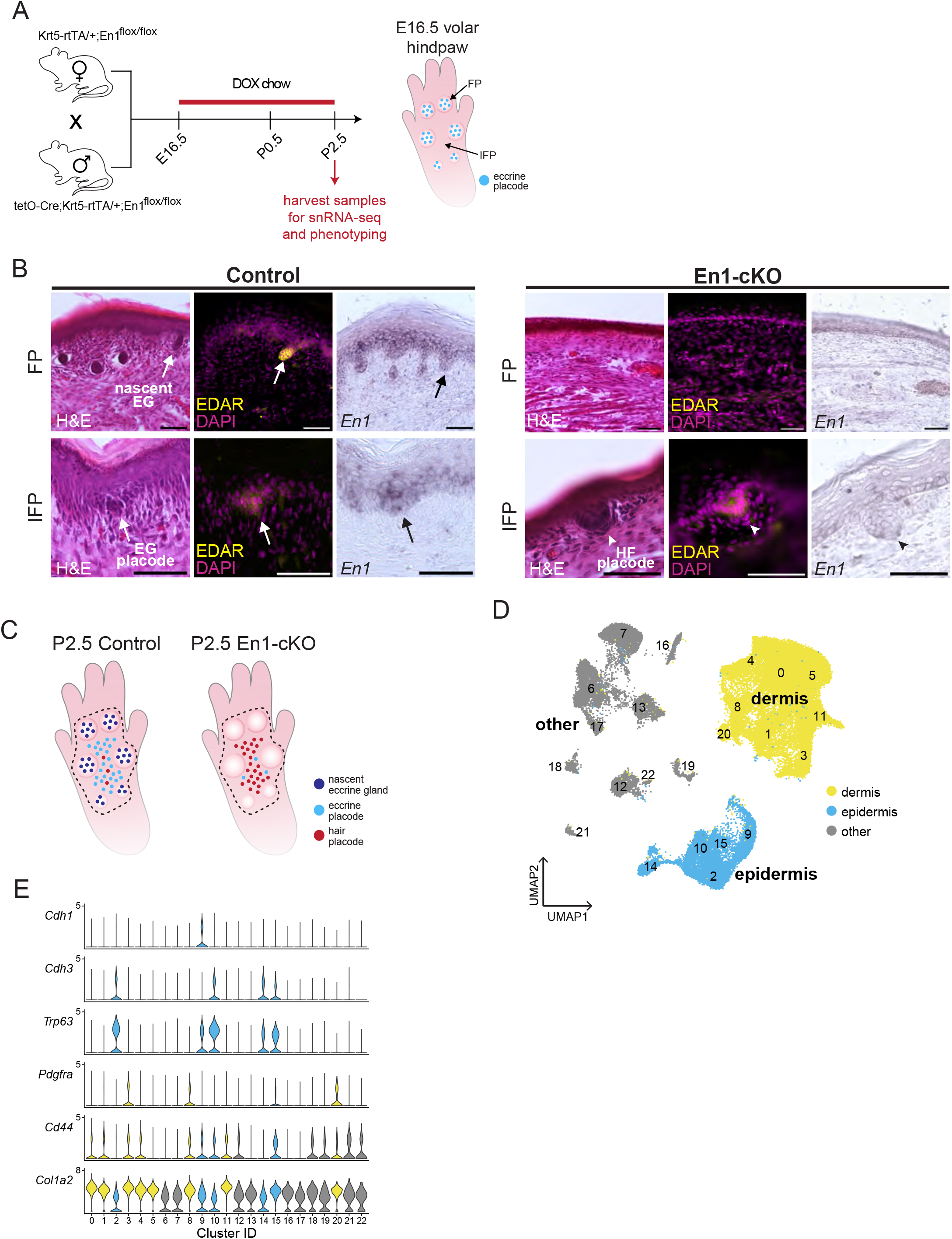
Single nucleus transcriptomic profiling of mouse skin after targeted disruption of eccrine placode identity. A) Mating scheme to generate animals, dosing schedule, and time points for analyses performed in this study. All En1-cKO and Control mice analyzed in this study were administered doxycycline (DOX) starting on embryonic day (E)16.5 until day of harvest. Volar hindpaw skin was harvested on post-natal day (P) 2.5 and analyzed as indicated. Schematic of E16.5 volar hindpaw shows eccrine placodes first form in the footpads. B) Representative images of stained adjacent sagittal sections through the footpad and interfootpad regions of P2.5 Control (left) and En1-cKO (right) volar skin. First section in series is stained with Hematoxylin and Eosin (H&E) for overall morphology; second section in series is stained with EDAR antibody (yellow) to visualize developing eccrine and hair appendages, as indicated. Section is counterstained with 4′,6-diamidino-2-phenylindole (DAPI; magenta). Hair placode associated dermal condensate (arrowheads). I*n situ* hybridization for *En1* (purple) on the third adjacent section in each series. Scale bars represent 50 µm. Eccrine gland (EG), Hair follicle (HF), Footpad (FP), Interfootpad (IFP). C) Schematic of Control (left) and En1-cKO (right) volar hindpaw skin at P2.5 summarizing where nascent glands (dark blue), eccrine placodes (light blue), and hair placodes (red) are found at this stage. Dashed line indicates the region of volar hindpaw skin that was dissected and snap frozen for snRNA-seq. D) UMAP projection of 45,370 nuclei from merged Control and En1-cKO datasets, which were analyzed in this study. Cluster identification numbers are annotated. Epidermal and dermal nuclei are highlighted in blue and yellow, respectively. All other nuclei are in gray. E) Violin plots of gene expression by cluster for markers of the epidermis (*Cdh1*, *Cdh3, Trp63*) and dermis (*Pdgfra*, *Cd44*, *Col1a2*).

Analyses of adult Control and En1-cKO mice confirm that the regimen we used to disrupt epidermal *En1* during development permanently inhibits eccrine formation (Figure S1A). We find that compared to Controls, the volar skin of adult En1-cKO mice exhibits a dramatic reduction in footpad eccrine glands coupled with a near-complete loss of these organs in the interfootpad space, as well as an increase in the number of fully formed, interfootpad hair follicles (Figure S1B-E; Tables S1–S3).

Our findings demonstrate that genetic manipulation of the *En1* locus in the basal epidermis can be used as a tool to specifically perturb the eccrine developmental program at its onset. Collectively, the reciprocal effect of *En1* on P2.5 eccrine and hair placode identities in the interfootpad space, and the loss of differentiating nascent eccrine glands in the footpads of En1-cKO mice at this stage, makes this an ideal experimental system to capture not only the eccrine developmental program but also the transitions that characterize its progression (Figure 1C).

### Single nucleus RNA-sequencing recovers transcriptional signatures of the primary skin layers

The volar hindpaw skin of P2.5 mice is composed of many different cell types, which have differential sensitivity to *En1* and varying degrees of involvement in eccrine gland development. Accordingly, we adapted Drop-seq-based, single nucleus RNA-sequencing (snRNA-seq) to recover the transcriptional profiles of P2.5 Control and En1-cKO skin samples^34–36^. Using the regimen described above, we dosed pregnant dams with doxycycline at E16.5 and collected the volar hindpaw skin of P2.5 Control and En1-cKO pups for snRNA-seq (Figure 1A, C). For each of the three profiled genotypes (two Control genotypes: En1^flox/flox^, Krt5-rtTA/+; En1^flox/flox^; one En1-cKO genotype: tetO-Cre/+;Krt5-rtTA/+;En1^flox/flox^), we profiled two biological replicates, where each replicate was comprised of pooled, volar hindpaw skins from five mice of the same genotype. In total, the transcriptomes of 45,370 sequenced nuclei passed quality control filtering and were interrogated in subsequent analyses.

After sample integration with Harmony, initial clustering on the merged data from all nuclei across Control and En1-cKO groups yielded 23 clusters (Figure 1D; Figure S2A)^37^. Analysis for marker gene enrichment reveals that our experiment captured the transcriptomes of nuclei from the two major compartments of the skin, namely nuclei of epidermal (blue) origin and of dermal (yellow) origin (Figure 1D, E; Figure S2B,C and Data S1). Characteristic of the epidermal skin layer, we find clusters that show enrichment for the epidermal makers *Cdh1* (e-cadherin), *Cdh3* (p-cadherin), and *Trp63* (p63) (9,070 nuclei; Figure 1E and Figure S2B, Data S1)^38–40^. The two Control genotypes contribute 6013 nuclei to the epidermal clusters, while the En1-cKO genotype contributes 3057 nuclei. Nuclei from clusters of a nominally dermal origin, which are the most abundant populations captured in our analyses, express known fibroblast markers including *Pdgfr alpha* (*Pdgrfa*), *Cd44*, and *Collagen type I alpha 2 (Col1a2*) (25,859 nuclei; Figure 1E, Figure S2C and Data S1)^41^. Of these, the two Control genotypes contribute 17,398 nuclei to the dermal clusters, and the En1-cKO genotype contributes 8461 nuclei. Based on marker enrichment, we additionally identify clusters of endothelial, smooth muscle, and immune origins (Figure 1D gray, Figure S2D, E and Data S1). These findings demonstrate that the nuclear transcriptome reflects the cellular heterogeneity of the volar skin and that our dataset captures expression profiles from the primary skin populations from both Control and En1-cKO experimental samples.

### A specialized transcriptome dominates the expression landscape of nascent stage eccrine glands

Like all ectodermal appendages, eccrine glands form from *Krt5* and *Krt14*-expressing basal keratinocytes^8–12^. All cells of the developing eccrine gland retain expression of these basal-specific epidermal markers from the placode stage through the nascent period^9, 11^. We observe enrichment for *Krt5* and *Krt14* in clusters 2, 10, 14, and 15, suggesting that the expression signatures of eccrine epidermal progenitors are likely to be contained within this subset of clusters (Figure 2A and Data S1). Consistent with this, we find that cluster 14 is almost entirely depleted of En1-cKO nuclei, accounting for just 1.64% of En1-cKO epidermal nuclei in the dataset but accounting for 17.71% of epidermal nuclei recovered from Controls (Figure 2B, C). *In situ* hybridization to detect *Trpv6* and other top marker genes of cluster 14, reveals a consistent pattern of enrichment in nascent eccrine glands in the hindpaw footpads of wildtype mice (Figure 2D, E; Figure S2A, B). That the nascent gland transcriptome is sufficiently diverged as to cluster separately from the other epidermal populations shows that at this stage of development, the overriding transcriptional character of eccrine epidermal progenitors is highly specialized and differentiated with respect to the rest of the epidermis.

**Figure 2.**
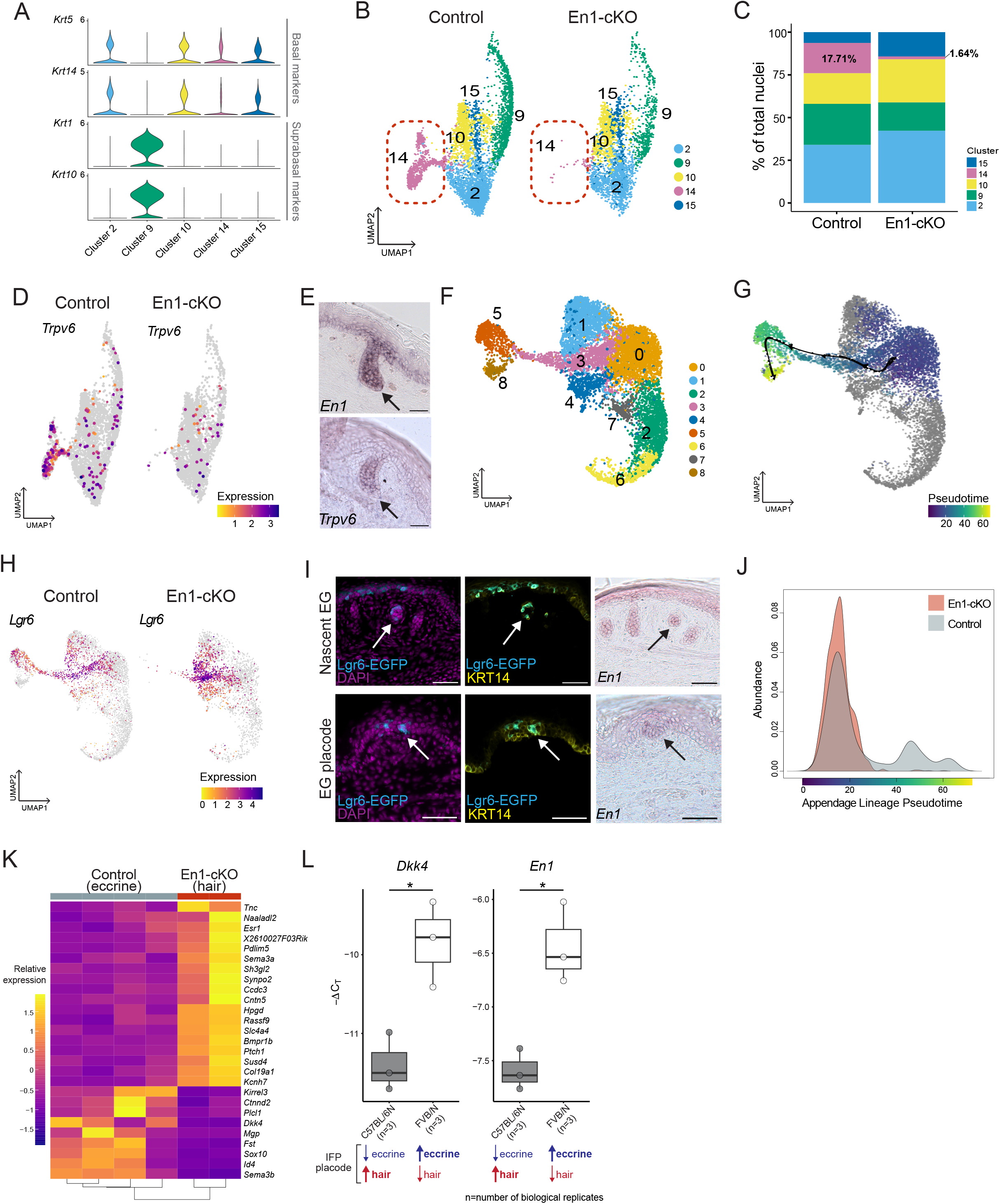
Transitions in the eccrine transcriptome during development. A) Violin plots of gene expression in only epidermal clusters for markers of basal keratinocytes (*Krt5*, *Krt14*) and suprabasal keratinocytes (*Krt1*, *Krt10*). B) UMAP projections of epidermal nuclei from original clustering (Figure 1D) colored by cluster identity and split by condition (Control and En1-cKO). Red dashed oval highlights cluster 14, which is depleted in En1-cKO. C) Percent contribution of nuclei from each epidermal cluster to total epidermal nuclei from Control and En1-cKO samples, respectively. Clusters are coded by color as depicted in B. D) Feature plots showing normalized expression of cluster 14 marker *Trpv6* in Control and En1-cKO epidermal nuclei. Negative nuclei are shown in gray. E) *In situ* hybridization for *En1* (top) and *Trpv6* (bottom) in a wildtype footpad at P2.5. Arrow indicates nascent eccrine gland. F) UMAP projection of subclustered epidermal nuclei colored by subcluster identity (merged Control and En1-cKO conditions). G) Results of slingshot trajectory inference mapped onto the subclustered epidermal UMAP embedding that depicts an inferred lineage through epidermal subclusters 0, 3, 5, and 8. Points representing nuclei are colored according pseudotime; nuclei not involved in this lineage are colored gray. H) Feature plot showing normalized expression of *Lgr6*, a top marker of epidermal subcluster 3, for Control (left) and En1-cKO (right) subclustered epidermal nuclei. I) Immunofluorescence staining of sagittal sections through the volar hindlimb skin of Lgr6-EGFP mice at P2.5. Tissue sections are stained with antibodies against EGFP (cyan), which reads out endogenous *Lgr6*, and Keratin 14 (KRT14; yellow) and 4′,6-diamidino-2-phenylindole (DAPI) nuclear stain (magenta). Sections through footpad skin containing nascent eccrine glands (top), and also of the interfootpad region that contains eccrine gland placodes (bottom, arrow) are shown. *In situ* hybridization for *En1* on adjacent serial sections is used to confirm eccrine gland (top and bottom, right). Scale bars represent 50 µm. J) Pseudotime density distribution of Control (gray) and En1-cKO (red) nuclei for slingshot lineage shown in E. Kolmogorov-Smirnov test of the distributions indicates differential progression along this trajectory between the conditions (p < 2.2e^-16^; D = 0.26487). K) Heatmap showing relative expression of transcripts that are differentially expressed between epidermal subcluster 3 nuclei from Control and En1-cKO samples, which are populated by eccrine glands and hair follicles, respectively (p adjusted < 0.01; absolute log2 fold change > 0.58). Columns are pseudobulk sample replicates and rows are genes. L) qRT-PCR results for *Dkk4* (left) and *En1* (right) of volar hindpaw skin from C57BL/6N (n=3) and FVB/N (n=3) mice at P2.5. The interfootpad space of C57BL/6N contains mostly hair placodes at P2.5, while that of FVB/N contains mostly eccrine placodes at this stage. Target gene expression is normalized to the reference gene *Rpl13a* and shown as -ΔC_T_. Significant differences between groups assessed by Mann-Whitney-Wilcoxon test (* p < 0.05).

### Identification of an eccrine developmental lineage in the epidermis

We specifically interrogated the heterogeneity of the epidermal clusters with a goal of resolving finer scale expression differences in the epidermis and identifying transcriptional signatures of pre-nascent stage eccrine gland progenitors including those of the eccrine placodes. After clustering only the epidermal nuclei isolated from merged Control and En1-cKO genotypes, we find that these further resolve into nine epidermal subclusters (Epi0-8) (Figure 2F; Data S2). Of these, Epi5 and Epi8 correspond to the validated nascent gland cluster 14 in the initial analysis, while nuclei in Epi2, Epi6, and Epi7 are characterized by enrichment for the suprabasal markers *Krt1* and *Krt10* (Figure S3C, D; Data S2)^9, 11^. Since early eccrine epidermal progenitors, including those of the placode, are basal in character, the nuclei within Epi0, Epi1, Epi3, or Epi4, which are enriched for the basal epidermal markers *Krt5* and *Krt14*, may represent these earlier stages of eccrine gland development (Figure S3C-E; Data S2).

Trajectory inference performed on all epidermal nuclei identifies a lineage that originates with cluster Epi0, which bears the transcriptional hallmarks of multi-potent basal epidermal keratinocytes, and ends with the validated nascent eccrine gland clusters Epi5 and Epi8 (Figure 2G; Figure S3D; Data S2). Epi3 represents an intermediate state along this inferred lineage and is enriched for transcripts involved in ectodermal appendage development including *Edar* (Figure S3D-F; Data S2).

To classify the Epi3 nuclei, we investigated the expression of *Lgr6*, which is the top upregulated transcript of this subcluster compared to all other epidermal populations (Figure 2H; Figure S3D; Data S2). *Lgr6* is known to be expressed in hair follicle placodes, suggesting that Epi3 may capture the transcriptional signatures of the earliest appendage primordia^56^. Consistent with this, we find that in the volar hindpaw skin of P2.5 *Lgr6-EGFP* knock-in reporter mice, EGFP is expressed in the eccrine placodes of the interfootpad space and in the nascent glands of the footpads (Figure 2H, I; Data S2). These collective data reveal that our snucRNA-seq dataset captures a continuous eccrine developmental lineage. Intriguingly, both Control and En1-cKO nuclei contribute to the early inferred eccrine lineage in the epidermis and indeed Control and En1-cKO samples cluster together in Epi0 and the early portion of Epi3, the origin and intermediate phases of the lineage, respectively (Figure 2J; Figure S3F). These observations suggest that during the initial stages of eccrine development, including during the placode stage, the transcriptional identity of eccrine progenitors is highly concordant with that of early hair follicle progenitors.

### An Engrailed 1-dependent signature of eccrine identity distinguishes the sweat gland placode from that of the hair follicle

Our findings reveal that nuclei derived from placode-stage volar appendages, both eccrine gland and hair follicle, are represented in Epi3. We reasoned that the transcriptional signature of the eccrine placode could be identified by specifically comparing the transcriptional content of Control (eccrine forming) versus En1-cKO (hair forming) nuclei within this intermediate cluster in the inferred eccrine epidermal lineage. We compared pseudo-bulk expression profiles of nuclei from Epi3 for Control versus En1-cKO samples (Figure 2K). This analysis identifies only 29 transcripts that are significantly differentially expressed between the Epi3 nuclei derived from the two conditions (Likelihood Ratio Test: adjusted p < 0.01; log2 fold change > 0.58). Of these, nine are relatively upregulated in the Control (eccrine) samples compared to the En1-cKO (hair) samples (Figure 2K).

Our targeted comparative analysis identified the secreted Wnt inhibitor *Dkk4* as a nuclear transcript that is relatively enriched in the eccrine containing Controls versus En1-cKO samples (Figure 2K). Notably, *Dkk4* is a known inhibitor of hair follicle formation, and *Dkk4* expression has previously been reported not only in hair follicle placodes but also in placode stage eccrine glands, which we confirmed by *in situ* hybridization in P2.5 volar skin of wildtype mice (Figure S3G)^10, 42–45^. To validate the quantitative enrichment of *Dkk4* in eccrine placodes relative to hair follicle placodes, we took advantage of strain specific differences in hair follicle and eccrine gland number in the hindpaw volar skin between C57BL/6N and FVB/N mice^19^. We have previously reported that, on average, the hindpaw interfootpad region of FVB/N mice contains 39 eccrine glands and 5 hair follicles, while the same region in C57BL/6N mice contains 8 eccrine glands and 65 hair follicles^19^. Using genetic mapping, we have previously implicated the higher expression of epidermal *En1* in FVB/N P2.5 volar hindpaw skin as the major causal driver for the specification of more eccrine gland placodes and fewer hair follicles placodes in this strain versus the P2.5 volar skin of C57BL/6N mice^19^. Analysis of *Dkk4* expression by qRT-PCR in P2.5 FVB/N and C57BL/6N volar hindpaws reveals a significant upregulation of *Dkk4* in the predominantly eccrine placode-containing skin of FVB/Ns relative to that of C57BL/6N (Figure 2L; Mann-Whitney-Wilcoxon test, *Dkk4*: p = 0.04953, Z = -1.964, r = -0.8). Notably, the effect size estimate of *Dkk4* upregulation is equivalent to that of *En1* upregulation in FVB/N as compared to C57BL/6N (Figure 2L; Mann-Whitney-Wilcoxon test, *En1*: p = 0.04953, Z = -1.964, r = -0.8).

The relative upregulation of *Dkk4* in eccrine gland versus hair follicle placodes confirms the existence of a specific signature of eccrine identity that is dependent on the epidermal expression of *En1*. Taking into account the extensive transcriptional similarity we observe for Control and En1-cKO nuclei early in the inferred eccrine epidermal lineage, our data are consistent with the concurrent but disproportionate representation of two distinct transcriptional programs in the eccrine placode: a major transcriptome of placode identity that is shared with the hair follicle; and a relatively underrepresented transcriptome that distinguishes the eccrine placode and is associated with its distinctive identity and unique biological properties.

### Identification of a transient, eccrine-associated dermal lineage

Skin-specific disruption of *En1* not only depletes eccrine-associated epidermal populations, but also results in dramatic depletion of En1-cKO nuclei from primary cluster 20, which is of dermal rather than epidermal origin (Figure 1D, Figure 3A). Nuclei from cluster 20 account for 1.99% and 0.08% of Control and En1-cKO dermal populations, respectively (Figure 3B). *S100a4* is the top marker gene that differentiates cluster 20 from all other dermal populations (Figure 3C; Figure S2C; DataS3). Using *in situ* hybridization, we find that *S100a4* is specifically expressed in a sheath of PDGFRA-positive dermal fibroblasts immediately surrounding the nascent stage eccrine glands in the footpads of P2.5 wildtype mice (Figure 3D). This pattern is recapitulated by *Tnc*, another top marker of cluster 20 (Figure S4A,B; Figure S2C; DataS3). The presence of a dermal sheath around nascent stage eccrine glands of mice was previously observed histologically by Cui and colleagues^10^. Consistent with the specific association of the cluster 20 transcriptional signature with dermal cells that surround the nascent glands, we find that *S100a4* expression is lost in the footpads of En1-cKO mice at P2.5 (Figure 3E).

**Figure 3.**
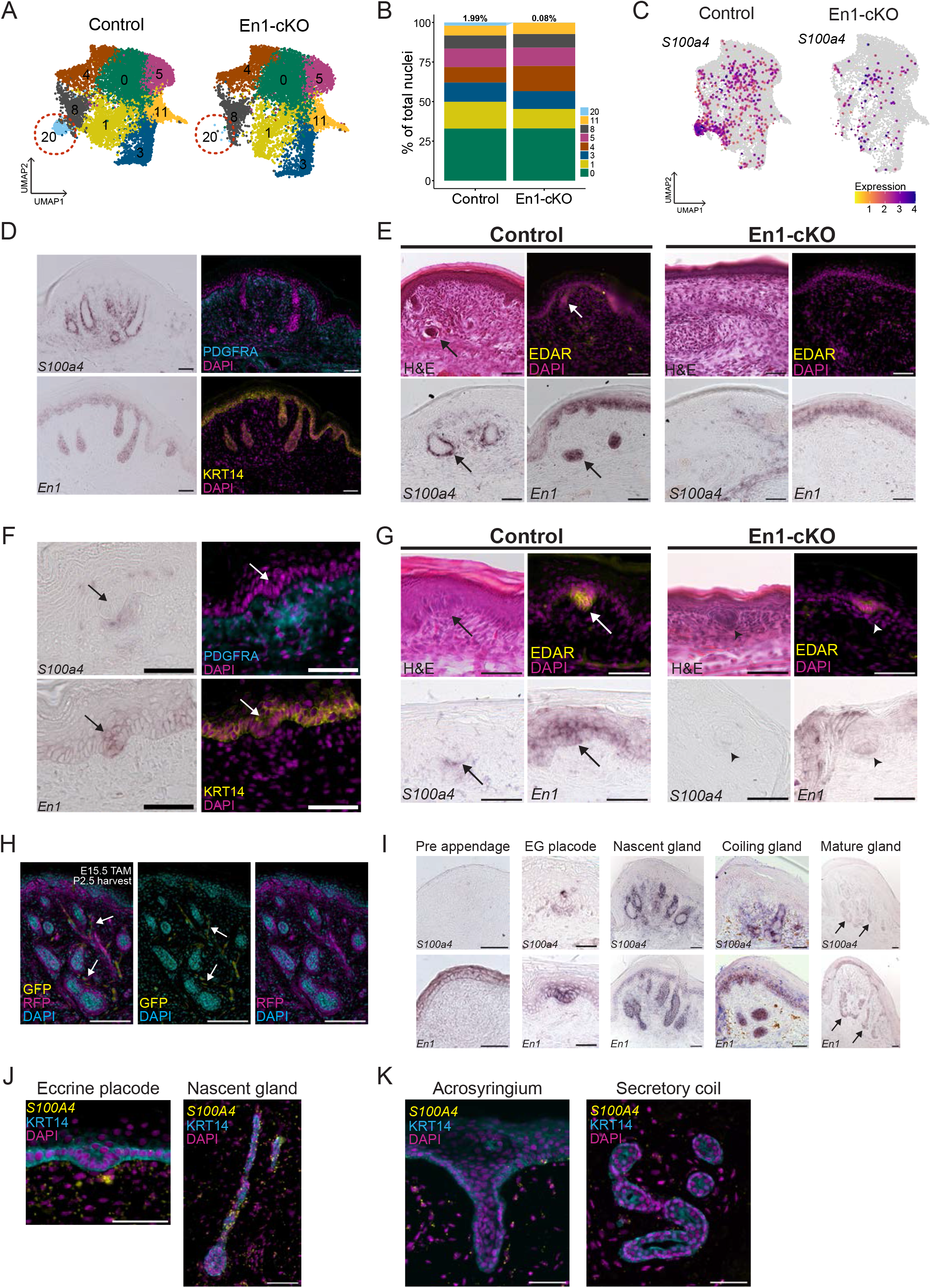
Identification of an *En1*-dependent dermal lineage associated with developing eccrine glands. A) UMAP projections of dermal nuclei colored by cluster identity (original clustering from Figure 1D) and split by condition (Control and En1-cKO). Red, dotted oval highlights cluster 20, which is depleted in En1-cKO. B) Percent contribution of nuclei from each dermal cluster to the total dermal nuclei from Control and En1-cKO samples, respectively. Subclusters are coded by color as depicted in A. C) Feature plots showing relative expression of cluster 20 marker *S100a4*, split by condition. D) *In situ* hybridization for *S100a4* (purple) in sagittal sections of wild type FVB/N P2.5 footpads containing nascent eccrine glands. *En1* expression (purple) by *in situ* hybridization is shown on adjacent sections. Immunofluorescence staining for PDGFRA (cyan), which marks dermal fibroblasts, and KRT14 (yellow), which marks basal keratinocytes. All images shown are from adjacent serial sections. E) Representative images of *in situ* hybridization for *S100a4* (purple) and *En1* (purple) along with EDAR immunofluorescence (yellow) and Hematoxylin and eosin (H&E) staining of adjacent serial sections of control (En1^flox/flox^) and En1-cKO (tetO-Cre; Krt5rtTA/+; En1^f/f^) footpad skin at P2.5. Arrows indicate nascent eccrine glands in the footpad. F) *In situ* hybridization for *S100a4* (purple) in sagittal sections of wild type FVB/N interfootpad skin containing eccrine placodes at P2.5. *En1* expression (purple) shown by *in situ* hybridization on adjacent sections. Immunofluorescence staining for PDGFRA (cyan) marks dermal fibroblasts and KRT14 (yellow) marks basal keratinocytes. All images shown are from adjacent serial sections. Arrows indicate eccrine placodes. G) Representative images of *in situ* hybridization for *S100a4* (purple) and *En1* (purple) along with EDAR immunofluorescence (yellow) and Hematoxylin and eosin (H&E) staining of adjacent serial sections of control (En1^flox/flox^) and En1-cKO (tetO-Cre; Krt5rtTA/+; En1^f/f^) interfootpad skin at P2.5. Arrows indicate eccrine placodes in the interfootpad region. Arrowheads indicate the hair placode and underlying dermal condensate in the interfootpad region of En1-cKO volar skin. H) Fluorescence images of a representative sagittal section through the footpad a S100a4-CreERT2/+;mTmG mouse at P2.5 that was given tamoxifen at E15.5 to label *S100a4*+ cells around placode stage in the footpad. S100a4-lineage cells (GFP) are shown in yellow. Membrane tdTomato (RFP), which marks all cells, is shown in magenta. Left image shows overlay of all channels. Middle image shows GFP (yellow) and DAPI (cyan) overlay. Right image shows RFP (magenta) and DAPI (cyan) overlay. I. *In situ* hybridization for *S100a4* (purple) and *En1* (purple) in volar skin at distinct stages of eccrine gland development: pre appendage (E15.5; footpad); eccrine placode (P2.5; interfootpad); nascent eccrine gland (P2.5; footpad); coiling eccrine gland (P7; footpad); mature eccrine gland (P28; footpad). J) RNAscope for *S100a4* (yellow) and immunofluorescence for KRT14 (cyan) in developing human plantar skin sections (120 gestational days) containing eccrine placodes (left) and nascent eccrine glands (right). K) RNAscope for *S100a4* (yellow) and immunofluorescence for KRT14 (cyan) in adult human cheek skin sections (64 years). Left image shows the mature eccrine gland acrosyringium and part of the duct and right image shows the secretory coil. All scale bars represent 50 µm (D-K). Nuclei are counterstained with DAPI in immunofluorescence images (magenta in D-G and J,K; cyan in H).

Intriguingly, we also observe *S100a4* expression specifically in the PDGFRA-positive dermal cells directly beneath eccrine placodes, which exhibit characteristic *En1* upregulation (Figure 3F, G). These *S100a4*-positive dermal cells are not morphologically distinct from the surrounding mesenchyme, consistent with previous reports that developing eccrine glands are never found in association with the dermal aggregates characteristic of the other major ectodermal appendages such as the hair follicle (Figure 3G)^8, 10, 21^. We find that the upregulation of *S100a4* in the eccrine placode-associated dermis is also dependent on epidermal *En1* since En1-cKO volar skin lacks dermal foci of *S100a4* including under the hair follicle placodes that populate the interfootpad space of this genotype (Figure 3G). By contrast, *S100a4* expression is retained in the ventral foot tendons of En1-cKO mice, indicating that its absence in the En1-cKO volar skin is specific to this context (Figure S4C). Using lineage tracing in S100a4-CreERT2/+; ROSA-mTmG mice, we find that the *S100a4*-expressing dermal cells under the eccrine placode give rise to the eccrine associated dermal sheath cells captured in cluster 20 (Figure 3H). We therefore conclude that epidermal *En1* expression is required for the induction of a single, eccrine-associated, dermal lineage.

*En1* expression in the limb ectoderm initiates well before the onset of eccrine gland formation in this tissue and persists in the adult, after development is completed^26, 29, 46^ (Figure 3I, bottom). In contrast, we find that *S100a4* expression in the dermis proximal to the eccrine glands is restricted to the developmental period during which the epidermis is actively forming these appendages(Figure 3I, top). Accordingly, we find no evidence of *S100a4* expression either prior to eccrine placode formation or around the mature eccrine glands of adult mice (Figure 3I; see also Figure S4D). This pattern of temporally restricted *S100a4* expression is recapitulated in human skin, in which we observe *S100a4* expression in the dermal fibroblasts associated with eccrine placodes and nascent glands (120 gestational days, plantar foot skin; Figure 3J) but not with any portion of the mature gland (64 years old, cheek skin; Figure 3K). Thus, the eccrine-associated dermal lineage is transient, appearing in conjunction with the activation of the *En1*-dependent epidermal eccrine developmental program in both mice and humans, and dissipating at its completion. Collectively, these data implicate, for the first time, an evolutionarily conserved and transcriptionally distinct dermal population that is specifically associated with developing eccrine glands.

### An *En1*-dependent eccrine niche (EDEN) is required for eccrine gland development

In light of finding that the *En1*-dependent dermal populations derive from a single lineage, and the specific association of these populations with developing eccrine glands in the epidermis, we interrogated the snRNA-seq data for evidence of signaling between the epidermal and dermal nuclei derived from Control samples. We first performed subclustering (Derm0-Derm11) and lineage pseudotime inference on all dermal nuclei to resolve putative developmental relationships within the broader dermal dataset (Figure 4A, B). We identify an inferred dermal lineage that terminates in subcluster Derm10, which corresponds to the validated, nascent gland-associated, dermal population represented in cluster 20 from the original analysis (Figure 3A, 4B). This lineage originates in Derm3, and successively progresses through Derm6, Derm9, and Derm2, before terminating in Derm10 (Figure 4B). Having identified the subset of dermal nuclei that constitute the putative eccrine-associated dermal lineage, we performed cell-cell interaction modeling between this subset of dermal clusters and those which make up the validated eccrine epidermal lineage (Epi0, Epi3, Epi5, and Epi8; Figure 2G) using CellChat^47^. We detect 185 significant receptor-ligand pairs from 32 pathways among the eccrine-associated dermal and epidermal subclusters (p < 0.05; Figure 4C; Figure S5A, B; Data S4). Of these, we identify 52 significant receptor-ligand interactions from 14 pathways that employ secreted signaling factors, including the WNT, BMP, TGFBeta, and non-canonical WNT (ncWNT) pathways (Figure S5A,B; Data S4). We identify receptor-ligand pairings consistent with bidirectional crosstalk between the lineages, with the epidermal lineage signaling to the dermal lineage and vice versa (Figure 4C; Figure S5A, B). Moreover, we find that the signals mediating these interactions change over developmental time (Figure 4C; Figure S5A, B).

**Figure 4.**
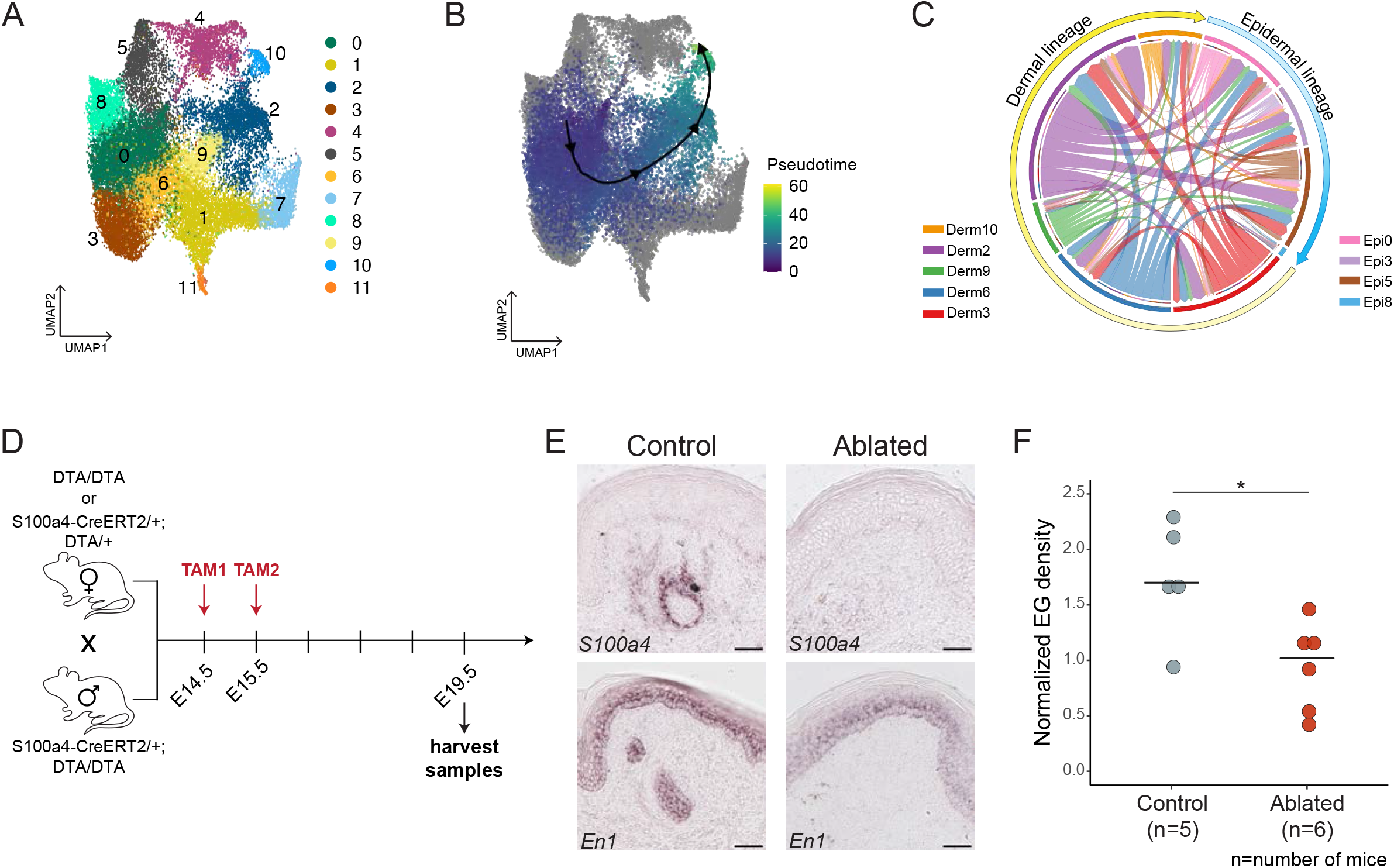
A transient *En1*-dependent eccrine niche (EDEN) is required for eccrine gland development. A) UMAP projection of subclustered dermal nuclei colored by subcluster identity (merged Control and En1-cKO conditions). B) Results of slingshot trajectory inference mapped onto the subclustered dermal UMAP embedding that depicts an inferred lineage through dermal subclusters 0, 3, 6, 9, 2 and 10. Points representing nuclei are colored according pseudotime; nuclei not involved in this lineage are colored gray. C) Summary chord diagram representing all significant signaling interactions between the dermal and epidermal lineage clusters inferred by CellChat (p < 0.05). Clusters are ordered around the circle based on their position in the inferred lineage. D) Experimental scheme for genetic ablation of *En1-*dependent dermal population during eccrine gland development. Tamoxifen-inducible S100a4-CreERT2 drives diphtheria toxin A (DTA)-mediated ablation of *S100a4* expressing cells. Pregnant dams were administered two doses of tamoxifen just prior to and overlapping with eccrine placode formation in the footpads (E14.5, E15.5) and pups were harvested via caesarean section at E19.5. E) Representative images of *in situ* hybridization for *S100a4* and *En1* from adjacent serial sections of an Ablated sample (S100a4-CreERT2/+; DTA/DTA) compared to a littermate Control (DTA/DTA). F) Dot plot of nascent eccrine gland density in the hindlimb footpads of Ablated and Control mice. Each dot represents the density of eccrine glands in an individual mouse, and one foot was analyzed per mouse. Eccrine gland number for each mouse was normalized to the number of sections scored for the analyzed foot. Cross bars represent sample medians. Significance was assessed by a Mann-Whitney U test (*p = 0.02846; Z = 2.1909; r = 0.66). All scale bars represent 50 µm.

The signaling interactions predicted from the snRNA-seq expression data suggest that the dermal and epidermal eccrine lineages engage in reciprocal crosstalk over the course of eccrine development. This finding is intriguing given the well-established importance of dermal-ectodermal interactions for the developmental progression of other major ectodermal appendages, such as the hair follicle and mammary gland^8^. Accordingly, we tested whether the newly identified *En1*-dependent dermal population is required for eccrine gland formation. For this purpose, we made use of the specific expression of *S100a4* in the mouse volar dermis to mark this population (Figure 3C-I; Data S3) .

Using an inducible genetic system, we ablated *S100a4*-expressing cells at the onset of eccrine gland development using targeted expression of diphtheria toxin subunit alpha (DTA) and assessed eccrine gland density in the volar skin of Control (S100a4-CreERT2/+;DTA/+ or DTA/DTA) as compared to Ablated (S100a4-CreERT2/+;DTA/DTA or S100a4-CreERT2/S100a4-CreERT2;DTA/DTA) mice (Figure 4D)^48^. In brief, timed pregnant females were dosed with tamoxifen to induce Cre activation at E14.5 and E15.5 to specifically ablate *S100a4* expressing cells around the time of footpad eccrine placode formation^9, 11, 12^. At E19.5, when footpad eccrine glands are at the nascent stage, we quantified eccrine gland density in the footpads of Control and Ablated mice (Figure 4D)^11^. This endpoint was selected due to dystocia in the dams, which necessitated end-point cesarean section to deliver pups for analysis.

We observe a deficit of *S100a4* positive cells in the footpads of Ablated genotypes as compared to Controls (Figure 4E). Notably, *En1* expression persists in both Control and Ablated groups (Figure 4E). The phenotypic significance of depleting the *S100a4* expressing dermal cells on eccrine gland development is striking. We find that eccrine gland density is on average 45% lower in Ablated compared to Control animals (Wilcoxon-Mann-Whitney test: p = 0.02846, Z = 2.1909, r = 0.66; Control: n = 5, Ablated: n = 6) (Figure 4E,F). Importantly, eccrine glands present in Ablated animals retained normal *S100a4* expression around the gland, indicating that the population was not effectively depleted in these regions (Figure S5C). Taken together, our findings identify an *En1*-dependent eccrine niche (EDEN) in the dermis, direct proximity to which is required for eccrine gland development in the ectoderm to proceed (Figure 5A).

**Figure 5.**
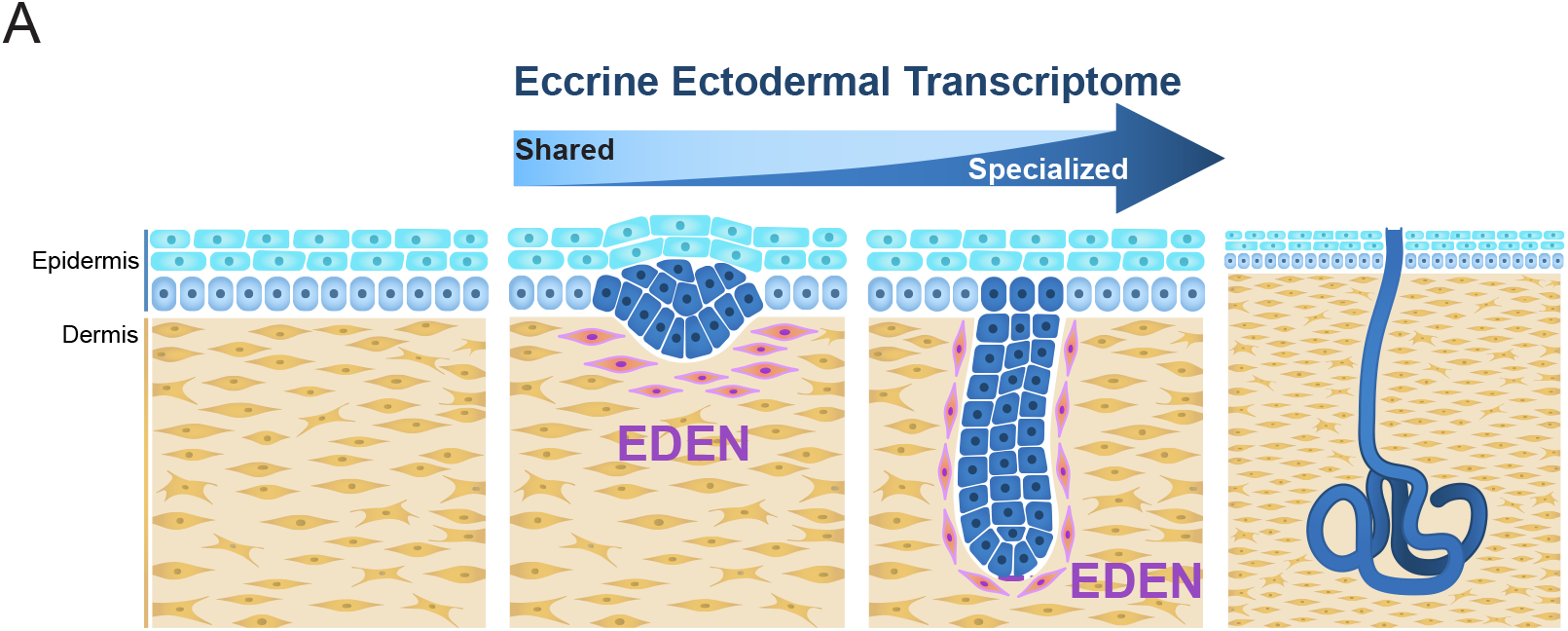
The transcriptional and cellular determinants of eccrine gland development. A) Model of eccrine gland development incorporating ectodermal and dermal determinants identified in this study. Eccrine gland development is coordinated by a progressive shift in the epidermal transcriptome from one that is predominantly shared with hair follicles, to a specialized, eccrine-specific transcriptome and requires the *Engrailed 1*-dependent eccrine niche (EDEN; pink) within the adjacent dermis. Epidermis is in blue, and dermis is shown in yellow. Pre- and inter-appendage basal epidermal keratinocytes are light blue, suprabasal keratinocytes are cyan, and eccrine cells are dark blue. Dermis is designated in yellow.

## DISCUSSION

In “The Variation of Animals and Plants Under Domestication”, Charles Darwin describes a family in Sindh (in present-day Pakistan) “in which ten men…were furnished, in the course of four generations…with only four small and weak incisor teeth and with eight posterior molars…have very little hair on the body, and … suffer much during hot weather from excessive dryness of the skin”^49^. One of the earliest formal accounts to suggest an underlying link between the formation of the group of organs we now call ectodermal appendages, Darwin’s description also aptly captures the importance of eccrine glands in human physiology. More than 150 years later, our study uncovers a specific blueprint for eccrine gland development and identifies transcriptional and cellular transitions at which the ectodermal appendage developmental paradigm diverges to build these essential organs (Figure 5A).

### Duality and temporal transitions in the eccrine gland epidermal program during development

We find that the upregulation of epidermal *En1* is necessary for a transcriptional signature that distinguishes the eccrine placode from that of the hair follicle. That this signature includes relative upregulation of *Dkk4*, a known inhibitor of folliculogenesis, suggests that developing eccrine glands may actively antagonize the formation of hair follicles^42, 43^. The upregulation of *Dkk4* is also significant because this gene encodes an inhibitor of canonical Wnt signaling, the level of which has been postulated to determine the fate of ectodermal appendage placodes, with hair placodes proposed to require the highest level of Wnt signaling among the different appendage types^12, 43, 50, 51^. Future studies are needed to distinguish between the eccrine placode-autonomous and – nonautonomous functions of *Dkk4*, and to understand the mechanisms by which these effects are mediated. Elucidation of how these functions are integrated with those of the other factors that constitute the *En1*-dependent transcriptome we identified in the eccrine placode sets the stage for uncovering the intrinsic program that confers eccrine identity in the epidermis.

The relative paucity of the specialized eccrine transcriptome as compared to the bulk of placodal transcripts that are shared between eccrine gland and hair follicle progenitors during early development contrasts with the expression profile of later stages. Our data indicate that as development proceeds, differentiation of the eccrine cell types is coupled with a concomitant shift in the balance of the eccrine transcriptome towards a more specialized and distinctive expression profile (Figure 5A).

Moreover, the extensive overlap we find in the transcriptome of hair and eccrine placodes may help to explain why previous efforts have also identified a largely shared set of signals in the early development of ectodermal appendages, irrespective of type^8, 10, 12, 14, 52, 53^. In particular, the relative dearth of specialized eccrine placode-enriched transcripts as compared to the far greater pool of those that are generalized across hair and eccrine types has intriguing implications for the character of the elusive, first dermal signal that triggers appendage fate and formation^12, 54–56^. Our observations raise the possibility that the first inductive signal from the dermis, which initiates eccrine development is also dualistic in nature. Given the relative proportions of the two classes of placodal transcripts captured in our study, the eccrine dermal signals may show a similar skew. This would in turn reduce the power to detect the specialized dermal inducers within the context of the greater proportion of those that are generalized across the different appendage types. The identification of the subset of transcripts that are differentially enriched in eccrine versus hair follicle placodes provides a directed means to parse out the precise composition of these upstream signals from the dermis.

### EDEN – a dermal niche for eccrine development

We find that the progression of the epidermal program in the eccrine placode requires the presence of EDEN, the first-described dermal niche for eccrine glands (Figure 5A). Importantly, our data show that EDEN is not the source of the first dermal inductive signal that initiates placode formation. That the transcriptional identity of EDEN depends on the upregulation of ectodermal *En1*, and that EDEN is found in association with eccrine gland epidermal anlagen, but not other *En1*-expressing cells, suggests that EDEN is analogous to the dermal condensations that form in response to the placodes of other major appendages such as teeth, hair, and mammary glands^8^. Future experiments to distinguish between the roles of *En1* in the placode as opposed to the inter-appendage epidermis in the induction of EDEN will help to clarify this. A starting point for these investigations and for functional dissection of the signals mediating interactions between the developing eccrine glands and the niche are the candidate factors identified in this study. Moreover, it is important to understand whether EDEN is required for eccrine development beyond the placode stage to establish its closest homology to the niches of other appendages. This is important because while the hair follicle dermal condensate (and its derivative the dermal papilla) is persistently required, the dense mesenchyme that directs early mammary gland development is transient and its role supplanted by the fat pad during the later stages of mammogenesis^8, 52^.

Irrespective of the precise source in the epidermis, our data demonstrate that a cell-autonomous transcriptional repressor, EN1, which is expressed in the basal epidermis, is required to induce a unique cellular identity in the underlying dermal compartment. There is precedence for this, as evidenced by the early and the separable role of ectodermal *En1* in maintaining the identity of the ventral limb bud by inhibiting mesenchymal *Lmx1b* expression^19, 26, 57^. In this context, *En1* acts by inhibiting *Wnt7a* in the ventral limb ectoderm^57^. Having identified a pool of *En1*-dependent transcripts in the eccrine placode and in the skin basal epidermis, our findings make it possible to determine whether *En1* effects are mediated by similar or entirely independent effectors in appendage development. Future studies to interrogate the function of these EN1 targets are also important to understand and distinguish between EN1’s roles in specifying the dermal niche, and in directly acting on the epidermal populations that make up the gland itself.

Intriguingly, we find that despite the continued expression of *En1* in adult basal keratinocytes in the eccrine forming regions and in the mature eccrine glands themselves, the transcriptional signature of EDEN does not persist at these stages in either mice or humans^29, 58^. This indicates that the dermal niche, at least as it exists during development, is not retained once eccrine gland formation is completed and may help to explain why eccrine glands exhibit limited ability to regenerate^9^. Previous studies have demonstrated the existence of several, unipotent stem cell populations within the mature eccrine gland, in the duct and the secretory coils, respectively^9^. In response to injury, these stem cells can only be mobilized to repair their respective lineages, but not the whole gland^9^. This contrasts with the regenerative ability of appendages such as the hair follicle, which retain multipotent regenerative ability and their dermal niche throughout life^59, 60^.

Importantly, the hair follicle dermal niche is not only necessary for the regeneration of the hair follicle but is also sufficient to induce epithelial cells to form *de novo* hair follicles, even in adulthood^60^. It is intriguing to speculate that EDEN, which our data indicate is the analogous dermal niche for the eccrine gland, may have comparable properties with respect to inducing *de novo* eccrine gland formation. Understanding whether the molecular effectors of EDEN demonstrate such eccrine-inducing capabilities has the potential to seed efforts to repair wounds and burns with skin regenerates that contain eccrine glands. Coupled with the capture of an eccrine epidermal transcriptome during development, the identification of EDEN not only addresses the long-standing developmental question of how the dermal and epidermal skin compartments of mammalian skin are differentially mobilized to build eccrine glands but also sets the stage for meeting a critical need in human regenerative medicine.

## ACKNOWLEDGMENTS

The authors thank Eric F. Joyce, Cliff Tabin, Pantelis Rompolas, Gabriella Rice, Sixia Huang, Paola Kuri, Stephanie Tsai, Elizabeth Grice, George Cotsarelis, Ed Morrissey, and Sarah Tishkoff for helpful discussions on this study. The authors thank Cliff Tabin, Marisa S. Bartolomei, Matthew Weitzman, Arjun Raj, and Klaus H. Kaestner for critical feedback on the manuscript. The authors thank Stacie Bumgarner for assistance with scientific illustration. We thank the Penn Skin Biology and Disease Resource Center (SBDRC) for use of Core A (P30-AR069589). HLD was supported by a National Institutes of Arthritis Musculoskeletal and Skin Diseases of the NIH (NIAMS) 5T32AR007465 award. SMM was supported by the Penn Academy for Skin Health and the SBDRC (P30-AR069589). HW was supported by a National Human Genome Research Institute of the NIH U01-HG012047 award. IAG was supported by a National Institute for Childhood Health and Disease of the NIH Award R24HD000836. YGK was supported by a SBDRC Pilot and Feasibility Award (P30-AR069589), a National Science Foundation (NSF) BCS-1847598 award, and a National Institute of Arthritis Musculoskeletal and Skin Diseases of the NIH Award R01AR077690. Any opinions, findings, and conclusions or recommendations expressed in this material are those of the authors and do not necessarily reflect the views of the NSF. The content is solely the responsibility of the authors and does not necessarily represent the official views of the NIH.

Birth Defects Research Laboratory (BDRL): Ian A. Glass^1^, Kimberly A. Aldinger^1, 2^, Dan Doherty^1^, Ian G. Phelps^1^, Jennifer C. Dempsey^1^, Kevin J. Lee^1^, Lucinda A. Cort^1^

1 University of Washington

2 Seattle Children’s Research Institute

## AUTHOR CONTRIBUTIONS

YGK conceived of and supervised the study. YGK and HLD designed the experiments and wrote the manuscript. HLD, RRT, AA, BK, PH, QQ, SMM, JCD, DA, and MM performed the experiments. HLD and YGK analyzed and interpreted the data. HW, IAG, the BDRL and YGK contributed resources and acquired funding to support the study.

## SUPPLEMENTAL FIGURE LEGENDS

**Figure S1.**
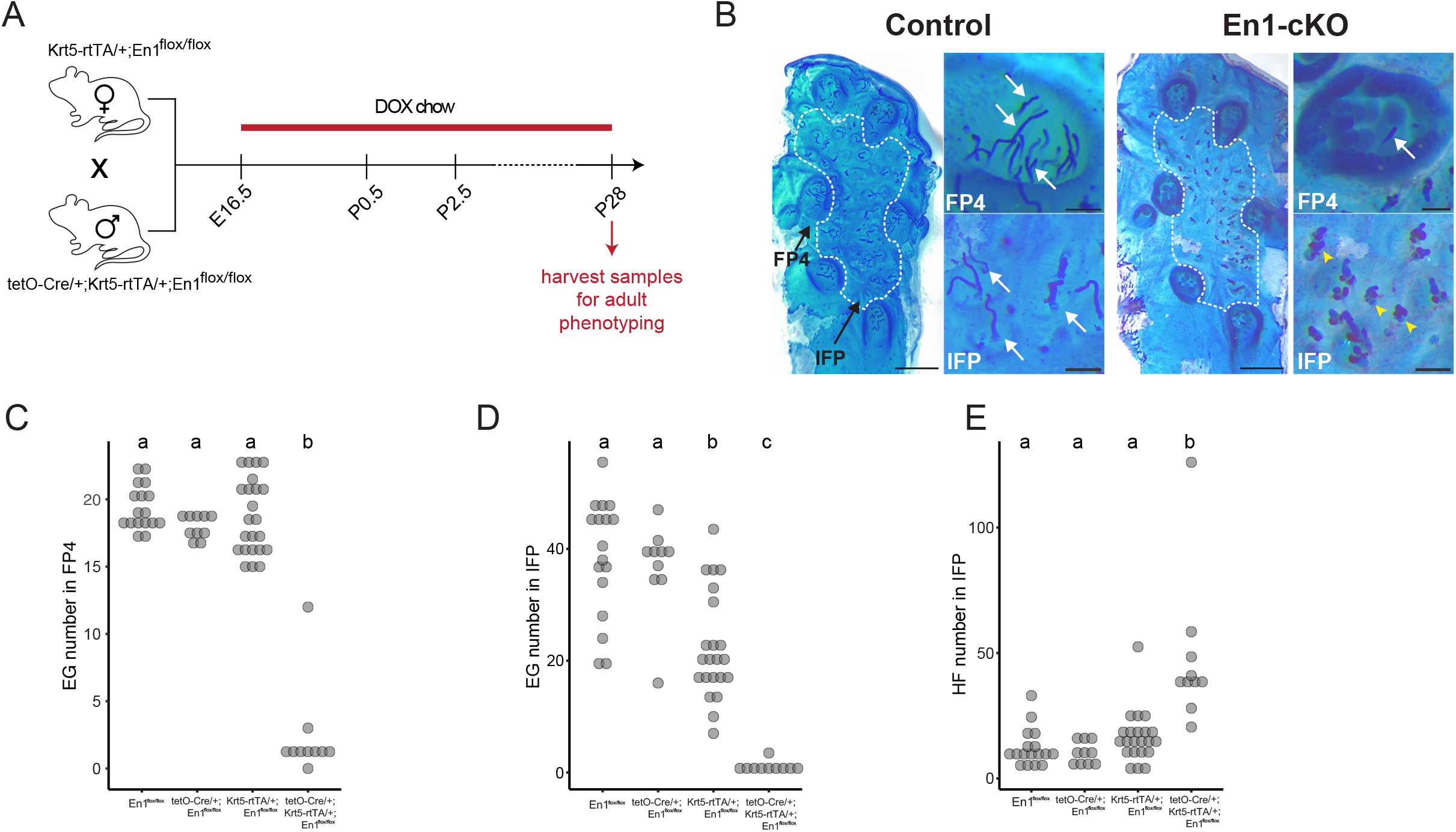
Appendage counts in Control vs. En1-cKO volar skin split by genotype. A) Mating scheme to generate animals, dosing schedule, and time points for analyses performed in this study. All En1-cKO and Control mice analyzed in this study were administered doxycycline (DOX) starting on embryonic day (E)16.5, when eccrine placodes first form in the footpads, until day of harvest. Volar hindpaw skin was harvested on post-natal day (P) 28 for adult phenotyping. B) Representative whole-mount preparations of P28 Control (left) and En1-cKO (right) hindlimb, volar skin. All appendages are stained with Nile blue (blue color) and hair follicle-associated sebaceous glands are stained with Oil Red O (red color). Footpad 4 is labeled and indicated by a black arrow. White dotted line outlines the interfootpad region (IFP). Scale bars in whole-mount images represent 1mm. Insets show higher magnification images of footpad 4 (FP4; top), and of the interfootpad regions (IFP bottom) of Control and En1-cKO skin. Eccrine gland (white arrow). Hair follicle (yellow arrowhead). Inset scale bars represent 200 µm. C) Plots of eccrine gland (EG) number in footpad 4 (FP4) showing samples split according to the specific genotypes that comprise Control and En1-cKO groups. D) Quantification of eccrine gland number in the interfootpad (IFP) from Figure 1E with samples split by genotypes that comprise Control and En1-cKO groups. E) Quantification of hair follicle (HF) number in the interfootpad region from Figure 1F with samples split by genotypes that comprise control and En1-cKO groups. Letter annotations on all plots (A-C) indicate significant pairwise differences found via Kruskal-Wallis followed by Dunn *post-hoc* testing (p<0.05). Groups annotated with the same letter are not significantly different from each other. Statistics reported in Tables S1–S3.

**Figure S2.**
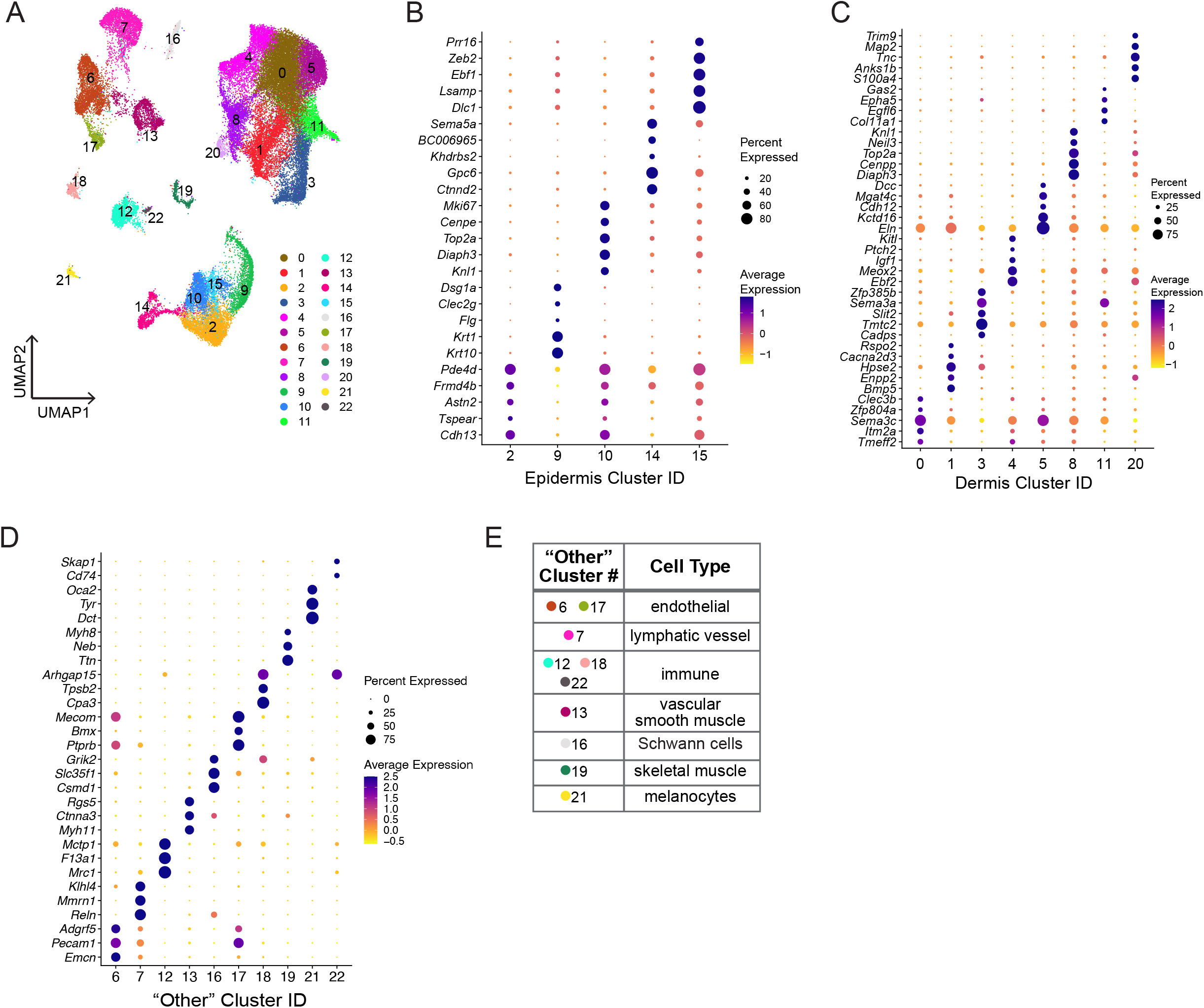
Clustering details for all nuclei from merged conditions. A) UMAP projection from main Figure 1D colored by cluster identity. B-D) Dot plots showing the expression of the top 5 marker genes identified for epidermal (B), dermal (C), and non-epidermal or dermal clusters (D; labeled “other” in main Figure 1). Dot color indicates average relative expression level for a cluster and dot size indicates the percent of nuclei in that cluster expressing the marker gene. E) Summary of cell type annotation for non-epidermal or dermal (“other”) clusters based on marker gene analysis.

**Figure S3.**
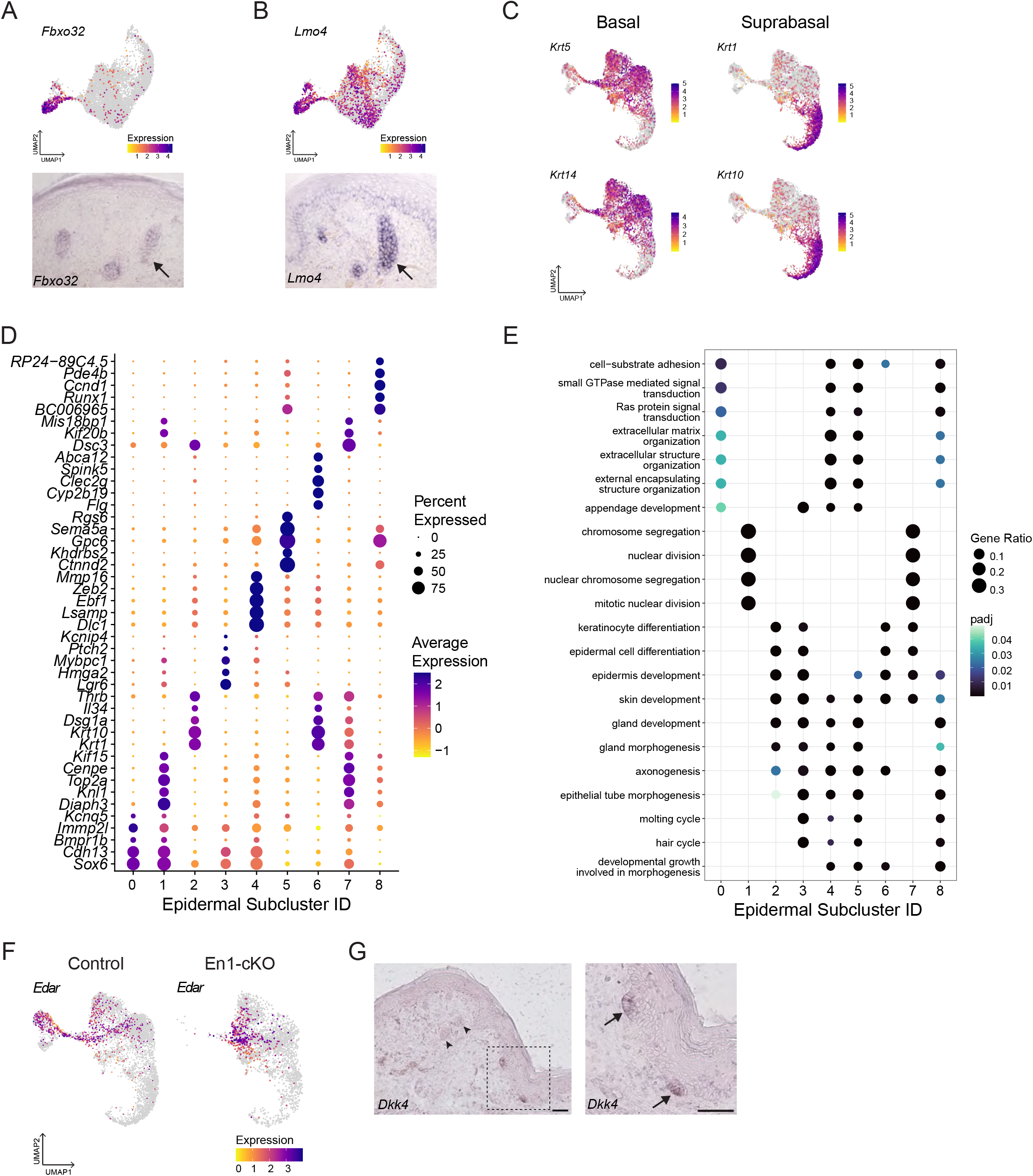
Epidermal subcluster marker genes and enrichment by gene ontology (GO) A) Feature plot (top) for *Fbxo32*, a specific marker gene of cluster 14. *In situ* hybridization for *Fbxo32* (bottom) in the volar limb in the footpad region at P2.5. B) Feature plot (top) for *Lmo4*, an additional marker of cluster 14. *In situ* hybridization for *Lmo4* (bottom) in the volar limb in the footpad region at P2.5. Arrows indicate nascent eccrine glands (A,B). C) Feature plots for known markers of basal keratinocytes (*Krt5*, *Krt14*) and suprabasal keratinocytes (*Krt1*, *Krt10*) showing normalized gene expression projected onto the UMAP embedding for subclustered epidermal nuclei. D) Dot plot depicting expression of the top 5 marker genes identified for each epidermal subcluster. Dot color indicates average relative expression level for the subcluster and dot size indicates the percent of nuclei in that subcluster expressing the marker gene. E) Dot plot depicting the top 4 results from a GO overrepresentation analysis conducted on significant differentially expressed marker genes for each epidermal subcluster (LFC ≥ 0.58, p adj. < 0.001). Dot color represents adjusted p value for GO term enrichment and dot size represents the gene ratio. F) Feature plot showing normalized expression of *Edar*, a known marker of ectodermal appendages, for Control (left) and En1-cKO (right) subclustered epidermal nuclei. G) *In situ* hybridization for *Dkk4* in wildtype FVB/N volar hindpaw skin at P2.5. Lower magnification image (left) includes both nascent eccrine glands in the footpad (*Dkk4* negative; arrowheads) as well as eccrine placodes at the edge of the interfootpad space (*Dkk4* positive). Image to right shows eccrine placodes, indicated by arrows, at higher magnification. Scale bars represent 50µm.

**Figure S4.**
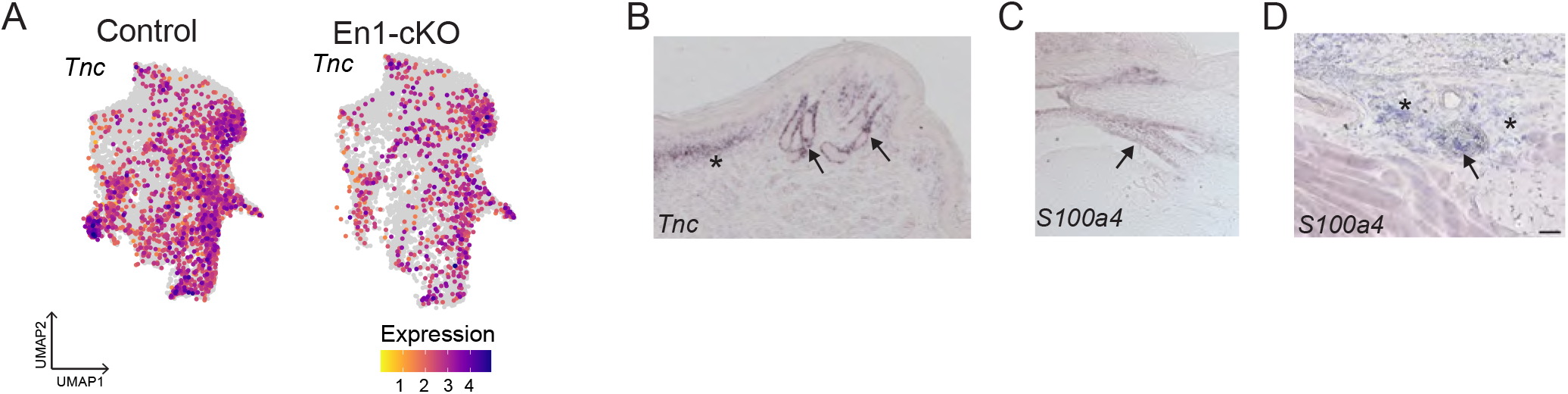
Additional validation of Cluster 20 markers. A) Feature plots show relative expression of *Tnc*, a marker gene of cluster 20, mapped onto UMAP projection of dermal nuclei from Control (left) and En1-cKO (right) samples. B) *In situ* hybridization for *Tnc* (purple) in the footpad region of the volar skin of a P2.5 FVB/N mouse. Nascent eccrine glands (arrows), as well as non-eccrine associated dermis (asterisk). C) *S100a4 in situ* hybridization signal (purple) in connective tissue of En1-cKO foot at P2.5 serves as internal positive control staining (same sample as shown in Figure 3E, right). D) *In situ* hybridization in the adult foot (P28) detects *S100a4* in a nerve (arrow) and connective tissue (asterisk) deep to the footpad. This serves as internal positive control for the mature eccrine gland *S100a4 in situ* shown in Figure 3I. Scale bars represent 50µm.

**Figure S5.**
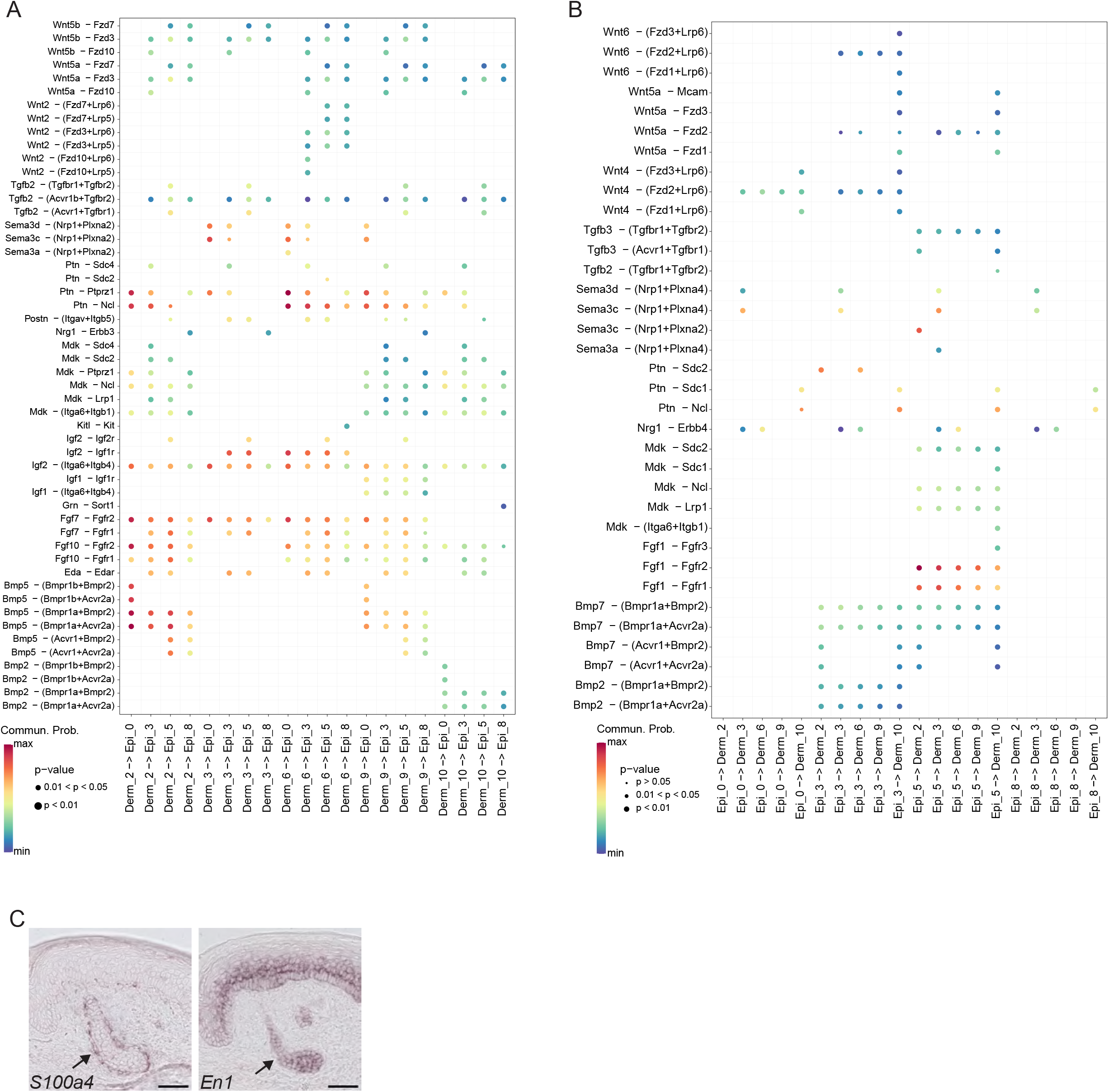
Inferred receptor-ligand interactions between dermal and epidermal subclusters and *S100a4* expression in volar skin. A,B) Bubble plots representing the communication probability for all significant receptor-ligand pair interactions inferred by CellChat (p < 0.05) showing signaling from dermis to epidermis (A) and epidermis to dermis (B). Dot color reflects communication probability and dot size indicates p value. C) *In situ* hybridization for *S100a4* (purple) and *En1* (purple) shows adjacent sections of Ablated volar skin with retained nascent glands. Sections shown are from the same Ablated sample as those shown in Figure 5C (bottom).

## METHODS

### Resource availability

#### Lead contact

Further information and requests for resources and reagents should be directed to and will be fulfilled by the corresponding author, Yana G. Kamberov.

#### Materials availability

This study did not generate new, unique reagents.

#### Data availability

Raw and processed snRNA-seq data files have been deposited in NCBI’s the Gene Expression Omnibus^61^ and are accessible through GEO Series accession number GSE220977 (https://www.ncbi.nlm.nih.gov/geo/query/acc.cgi?acc=GSE220977). Any additional information required to reanalyze the data reported in this paper is available from the lead contact upon request.

### Experimental model and subject details

#### Animals

*Tg(tetO-cre)1Jaw/J (tetO-Cre)*^32^ and *Gt(ROSA)26Sor^tm^*^1^(DTA)*^Lky^/J* (ROSA-DTA)^48^ mice were purchased from the Jackson Laboratory. *Tg(KRT5-rtTA)T2D6Sgkd/J* (Krt5-rtTA)^33^ mice were provided by Dr. Sarah Millar, *En1tm8.1Alj/J* (En1^flox^)^31^ mice were provided by Dr. Alexandra Joyner and *Lgr6tm2.1(cre/ERT2)Cle/J* (Lgr6-eGFP)^62^ mice were provided by Dr. Pantelis Rompolas. S100a4-CreERT2 mice were generated by Dr. Mayumi Ito and provided under material transfer agreement.

*Gt(ROSA)26Sortm4(ACTB-tdTomato,-EGFP)Luo/J* (mTmG)^63^ mice were provided by Dr. George Cotsarelis. En1^flox^, Krt5-rtTA, and tetO-Cre mice were maintained on an FVB/N background (Charles River Laboratories) for more than 10 generations. ROSA-DTA mice were maintained on a C57BL/6J background (Jackson Laboratory) until they were bred to S100a4-CreERT2 mice, which were maintained on a mixed genetic background. Lgr6-eGFP mice were also maintained on a mixed genetic background. FVB/N and C57BL/6N wild type mice were obtained from Charles River Laboratories and Taconic Biosciences, respectively. Both males and females were used for all experiments in this study.

For inducible En1 conditional knock-out (En1-cKO) experiments, male mice harboring the tetO-Cre, Krt5-rtTA, and homozygous En1^flox^ alleles were mated to females homozygous for En1^flox^ and carrying the Krt5-rtTA allele. Timed pregnant dams were continuously administered a doxycycline dosed diet (6 g/kg) *ad libidum* beginning at E16.5. Offspring were euthanized at either P2.5 or P28 via decapitation or CO_2_ inhalation, respectively.

For S100a4 lineage tracing experiments, male S100a4-CreERT2/+;mTmG/+ mice were mated to mTmG/+ or mTmG/mTmG females. Timed pregnant females were dosed with tamoxifen in corn oil via oral gavage (3 mg/kg) at E15.5 and pups were euthanized at P2.5 for volar hindpaw collection.

For S100a4+ cell ablation experiments, male mice harboring at least one copy of the S100a4-CreERT2 and ROSA-DTA alleles were mated to females either homozygous for ROSA-DTA or heterozygous for both S100a4-CreERT2 and ROSA-DTA. Timed pregnant females were dosed with tamoxifen in corn oil via oral gavage (3 mg/kg) at both E14.5 and E15.5. Pregnant dams were euthanized at E19.5 (developmental equivalent of P0) and pups were harvested via endpoint caesarean section due to dystocia in the dams.

All mice used in this study were housed on a 12h light/dark cycle in a University Laboratory Animal Resources (ULAR) managed vivarium at the Perelman School of Medicine (PSOM). All procedures were performed in accordance with guidelines established by the National Institutes of Health and have been approved by the University of Pennsylvania PSOM Institutional Animal Care and Use Committee (protocol # 806105).

#### Human tissue

Specimens from fetal (120 gestational days) human plantar foot skin were obtained from the Birth Defects Research Laboratory at the University of Washington with ethics board approval and maternal written consent. Deidentified samples from adult human cheek skin (64 years old) were obtained from the Skin Biology and Disease Resource Center at the University of Pennsylvania with ethics board approval and written consent. This study was performed in accordance with ethical and legal guidelines of the University of Pennsylvania institutional review board.

### Method details

#### Quantification of appendages in adult volar skin

Quantification of eccrine glands and hair follicles at P28 was performed in whole mount preparations of hindlimb volar skin as previously described^16^. Briefly, the epidermis was separated from the underlying dermis by dispase digestion and fixed with 10% neutral buffered formalin. Hair follicle-associated sebaceous glands were stained with Oil Red O and appendages (eccrine glands and hair follicles) were stained with Nile Blue, and whole mounts were imaged on a Leica M680 dissection scope fitted with a Leica IC90E camera. All data reported in this manuscript represents the average counts from the right and left hindlimb volar skin of an individual mouse.

#### Isolation and purification of nuclei for snRNA-seq

En1-cKO and Control pups were euthanized at P2.5 and volar hindlimb skin (right and left) was rapidly dissected on ice and snap frozen on dry ice and stored at – 80°C until nuclear isolation. Single nuclear suspensions were generated from the pooled volar hindlimb skin (right and left) from 5 mice of the same genotype, for a total of 10 pooled volar skins, as previously described with some modifications^35^. Pooled samples were dissociated in homogenization buffer using a Polytron homogenizer (Kinematica Inc.) followed by dounce homogenization (20 times with loose pestle; 10 times with tight pestle). Suspensions were filtered to remove debris and single nuclear suspensions were quantified via hemocytometer after staining with Trypan Blue, which stains all nuclei. For P2.5 volar skin, we obtained ≥ 2.1x10^5^ nuclei/ml for each sample. Sucrose gradient centrifugation was not performed on single nuclear suspensions. Biological replicates were prepared and sequenced in two separate batches; one replicate of each genotype (En1-cKO: tetO-Cre/+; Krt5-rtTA/+; En1^flox/flox^; Control: En1^flox/flox^, Krt5-rtTA/+; En1^flox/flox^) was processed and sequenced in the same batch to avoid autocorrelation of condition and batch effects.

#### snRNA-seq library preparation and sequencing

Single nuclei were co-encapsulated with barcoded beads (ChemGenes) and reverse transcription was performed as previously described^35^. Optimal PCR cycle number for library amplification was determined via qPCR using 6,000 beads, cDNA was tagmented, and libraries were further amplified as previously described^35^. cDNA libraries were quantified via Qubit 3.0 (Invitrogen) and library quality was determined via Bioanalyzer (Agilent) prior to sequencing on an Illumina NextSeq 500 using the 75-cycle High Output v2 kit (Illumina). In total, six neonatal volar skin snRNA-seq libraries were sequenced in four sequencing runs. See open access snRNA-seq protocol for more details: https://www.protocols.io/view/snucdrop-seq-protocol-n2bvjr36plk5/v235.

#### *In situ* hybridization, immunofluorescence, RNAscope, and imaging

Volar skin was fixed in 4% PFA, cryoprotected using sucrose, and embedded in OCT (Tissue Tek) for cryo-sectioning at a thickness of 10-12 µm. Sections were collected in 3-4 series for staining of adjacent sections. *In situ* hybridization was performed as previously described^19^ with custom anti-sense DIG-labeled riboprobes transcribed *in vitro* (Roche) from amplified cDNA (see Key Resources Table for primers). Immunofluorescence was performed for KRT14 (1:10,000, BioLegend 905303), EDAR (1:100, R&D Systems AF745), PDGFRA (1:100, R&D Systems AF1062), and GFP (1:1000, Abcam ab13970) after blocking tissue in PBT (0.2% Triton-X100 in PBS) with 10% serum. Secondary detection was performed with antibodies conjugated to Alexa Fluor^488^ (Invitrogen A-11039 or Abcam ab150129), Alexa Fluor^594^ (Jackson ImmunoResearch 711-585-152), or Alexa Fluor^647^ (Jackson ImmunoResearch 711-605-152) and samples were counterstained with 4′6-diamidino-2-phenylindole (DAPI, Sigma D9542).

FFPE adult and fetal human tissue was processed and sectioned by SBDRC Core A. RNAscope was performed on 5 µm paraffin sections after standard target retrieval for *S100A4* (ACD Bio #422071)*, PPIB* (positive control; ACD Bio #313901), and *DapB* (negative control; ACD Bio #310043) using an RNAscope 2.5 HD-RED assay kit (ACD Bio #322350). Subsequently, immunofluorescence for KRT14 was performed as described above and samples were counterstained with DAPI.

Samples were imaged on a Leica DM5500B microscope equipped with Leica DFC 500 (bright field) and Leica DFC 360X (fluorescence) cameras. Images were processed using FIJI software^64^.

#### Scoring of footpad eccrine glands at E19.5

*S100a4* Ablated and Control samples were harvested as described above and whole hindlimbs were cryo-sectioned at 10 µm and three adjacent series of sections were collected. The first series was stained with hematoxylin and eosin (H&E, Sigma Aldrich) for scoring. The second and third series were used for *in situ* hybridization for *En1* and *S100a4*, respectively, to confirm efficient ablation of *S100a4*+ cells. Scoring of E19.5 Hematoxylin and eosin-stained sections was performed by counting all nascent eccrine glands in the footpads throughout the entire series of sections. Eccrine counts were then normalized to the number of sections scored to yield an eccrine density estimate.

#### RNA extraction and qRT-PCR

FVB/N and C57BL6/N neonatal pups were euthanized at P2.5 and volar hindpaw skin was harvested and snap frozen as described above. Biological replicates are comprised of left and right hindpaw skin pooled from 3 mice. Pooled skin samples were dissociated using a Polytron homogenizer and total RNA was isolated using TRIzol (Life Technologies) extraction followed by clean up and on-column DNase I treatment using an RNeasy Mini Kit (Qiagen). cDNA was reverse transcribed using SuperScript III (Thermo Fisher) following the manufacturer’s instructions. qRT-PCR was performed in biological triplicate and technical quadruplicate using Power SYBR PCR master mix (Thermo Fisher) for *Dkk4* and *En1*. *Rpl13a* served as a reference gene for normalizing Ct values. Each data point reported in Figure 2L represents the mean of technical replicates for a sample. Primers used for qRT-PCR are found in Table S4.

### Quantification and statistical analysis

#### snRNA-seq data pre-processing

Paired-end snRNA-seq reads were processed using publicly available the Drop-seq Tools v1.12 software^65^ with modifications described previously^35^. A digital expression matrix was generated by assembling a list of UMIs in each gene (as rows) within each cell (as columns), and UMIs within ED = 1 were merged.

#### snRNA-seq cluster identification and marker gene analysis

Digital expression matrices for each sample were loaded into Seurat v4.0.2^66^ and merged. UMI counts were normalized by scaling by library size, multiplying by 10,000, and transforming to log scale. Genes were filtered out if they were expressed in <10 nuclei and nuclei were filtered out if they contained a high proportion of UMIs mapped to mitochondrial genes (≥0.05), fewer than 300 or more than 6000 detected genes, resulting in 45,370 nuclei in the final filtered dataset.

Prior to clustering, individual sample datasets were integrated using the Harmony algorithm^37^, the top 2000 variable genes were identified using the *FindVariableFeatures* function in Seurat with the VST selection method, and expression of these variable genes was scaled and centered for principal components analysis. Based on the cumulative standard deviations of each principal component (PC), visualized by the function *ElbowPlot* in Seurat, we selected the first 40 PCs for two-dimensional uniform manifold approximation and projection (UMAP) implemented by Seurat *RunUMAP* with default parameters. Initial clustering on the full dataset (all nuclei, merged conditions) identified 23 clusters with the resolution parameter for *FindClusters* set to 0.7. Based on the expression pattern of well-established marker genes in this dataset, we assigned 25,859 nuclei to dermal cells (∼57% of our data), 9,070 nuclei to epidermal cells (∼20% of our data), and 10,441 nuclei to various other cell types (see Figure S2). Differential expression analysis was performed to identify marker genes for each of these 23 initial clusters using Seurat’s *FindAllMarkers* function with Wilcoxon test. Differentially upregulated genes that were expressed in at least 25% of nuclei with a log fold change (LFC) ≥ 0.58 were considered marker genes (Data S1). Dermal cluster specific marker genes were identified using the same approach, but with only dermal clusters of nuclei compared to each other (Data S3).

#### Subclustering, pseudobulk differential expression, trajectory inference, and CellChat analysis

Nuclei from clusters determined to be epidermal (original clusters 2, 9,10,14, and 15) or dermal (original clusters 0, 1, 3, 4, 5, 8, 11, 20) were subsetted, the top 2000 variable genes were identified, and PCA was performed as above. The top 40 PCs were selected for clustering analysis with a resolution of 0.4 for epidermal nuclei and 0.7 for dermal nuclei, leading to the identification of 9 epidermal subclusters and 11 dermal subclusters. Of these 9 epidermal subclusters, 6 are basal (6,799 nuclei) and 3 of these subclusters are suprabasal (2,271 nuclei). Marker genes of each subcluster were identified as described above (Data S2, S3) and gene ontology (GO) enrichment analysis was performed on these marker gene lists using the *compareCluster* function from the Cluster Profiler R package^67^.

We performed pseudo-bulk analysis to identify differentially expressed transcripts between Control and En1-cKO placode nuclei (Epi3). Pseudobulk samples were created using the Seurat function *PseudobulkExpression* on sample counts for each subcluster using the aggregate pseudo-bulk method. Differential expression analysis was performed on these pseudo-bulk samples for Epi3 using DESeq2^68^ with a likelihood ratio test. Genes were deemed significantly differentially expressed between Control and En1-cKO pseudo-bulk samples if p adjusted < 0.01 and LFC ≥ 0.58.

Trajectory inference of the subclustered nuclei was performed using Slingshot^69^ on either the subsetted epidermal or dermal PCA reduction (merged conditions). Epi0 was supplied as the starting cluster to the *slingshot* function for the epidermal trajectory inference; for dermal trajectory inference, subcluster Derm11 was provided as the end cluster. The inferred lineages were then mapped onto UMAP embeddings for visualization using the *embedCurves* Slingshot function. Differential progression analysis was performed on the epidermal lineage via Kolmogorov-Smirnov test of the lineage pseudotime distributions between Control and En1-cKO nuclei.

CellChat^47^ was performed on Control nuclei from the dermal and epidermal subclusters that were determined to be involved in the relevant inferred lineages using the default CellChat mouse database. Average gene expression per cluster was computed using the “truncatedMean” method (trim=0.1) within CellChat’s *computeCommunProb* function. All significant receptor-ligand interactions (p < 0.05) identified between any pair of clusters are summarized as a chord diagram (Figure 4C). Complete results are provided as a supplemental data file (Data S4).

#### Statistical analysis of phenotypic data and qRT-PCR expression

Statistical parameters, including sample sizes, are reported in figures, figure legends, tables, and/or corresponding results text. A Kruskal-Wallis test followed by a *post-hoc* Dunn’s test with Benjamini-Hochberg correction for multiple comparisons was performed on P28 appendage counts for En1-cKO phenotyping (alpha = 0.05). A Mann-Whitney-Wilcoxon test was used for pairwise comparisons of qRT-PCR data (alpha = 0.05). All statistical analyses were performed in R^70^.

## SUPPLEMENTAL TABLE LEGENDS

**Table S1.** Results of statistical analysis of the effect of genotype on footpad 4 (FP4) eccrine gland number in adult mice. Results of Kruskal-Wallis test followed by *post-hoc* Dunn’s test with Benjamini-Hochberg (BH) correction related to Figure S1A. Genotype codes: “floxCtrl” refers to En1^flox/flox^; “creCtrl” refers to tetO-Cre/+; En1^flox/flox^; “rttaCtrl” refers to Krt5-rtTA/+; En1^flox/flox^: “ko” refers to tetO-Cre/+;Krt5-rtTA/+;En1^flox/flox^. Alpha = 0.05.

| **Kruskal-Wallis FP4 ~ genotype** |  |  |  |
| --- | --- | --- | --- |
| chi-squared = 26.869, df = 3, p-value = 6.271e-06 |  |  |  |
| **FP EG Dunn test results** |  |  |  |
| Comparison | Z | p.unadj | p.adj (BH) |
| creCtrl - floxCtrl | -1.38 | 0.17 | 0.25 |
| creCtrl - ko | 3.23 | 1.25E-03 | 2.49E-03 |
| floxCtrl - ko | 5.00 | 5.77E-07 | 3.46E-06 |
| creCtrl - rttaCtrl | -0.53 | 0.60 | 0.60 |
| floxCtrl - rttaCtrl | 1.09 | 0.28 | 0.33 |
| ko - rttaCtrl | -4.34 | 1.43E-05 | 4.30E-05 |

**Table S2.** Results of statistical analysis of the effect of genotype on interfootpad (IFP) eccrine gland (EG) number in adult mice. Results of Kruskal-Wallis test followed by *post-hoc* Dunn’s test with Benjamini-Hochberg (BH) correction related to Figure S1B. Genotype codes: “floxCtrl” refers to En1^flox/flox^; “creCtrl” refers to tetO-Cre/+; En1^flox/flox^; “rttaCtrl” refers to Krt5-rtTA/+; En1^flox/flox^: “ko” refers to tetO-Cre/+;Krt5-rtTA/+; En1^flox/flox^. Alpha = 0.05.

| **Kruskal-Wallis IFP EG ~ genotype** |  |  |  |
| --- | --- | --- | --- |
| chi-squared = 38.202, df = 3, p-value = 2.561e-08 |  |  |  |
| **IFP EG Dunn test results** |  |  |  |
| Comparison | Z | p.unadj | p.adj (BH) |
| creCtrl - floxCtrl | -0.41 | 0.68 | 0.68 |
| creCtrl - ko | 4.67 | 3.00E-06 | 9.00E-06 |
| floxCtrl - ko | 5.65 | 1.56E-08 | 9.36E-08 |
| creCtrl - rttaCtrl | 2.49 | 1.26E-02 | 1.51E-02 |
| floxCtrl - rttaCtrl | 3.46 | 5.48E-04 | 1.10E-03 |
| ko - rttaCtrl | -2.98 | 2.86E-03 | 4.29E-03 |

**Table S3.**
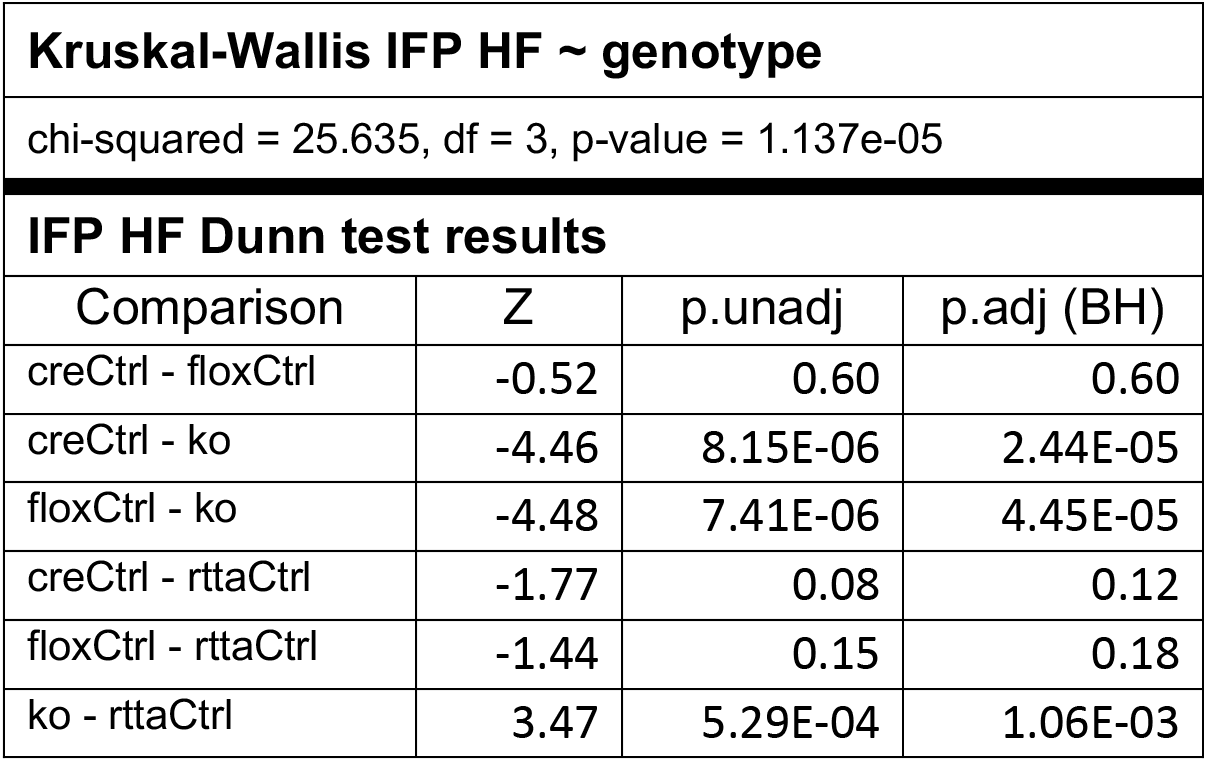
Results of Kruskal-Wallis followed by post-hoc Dunn’s test on the effect of genotype on interfootpad (IFP) hair follicle (HF) number in adult mice. Results of Kruskal-Wallis test followed by *post-hoc* Dunn’s test with Benjamini-Hochberg (BH) correction related to Figure S1C. Genotype codes: “floxCtrl” refers to En1^flox/flox^; “creCtrl” refers to tetO-Cre/+; En1^flox/flox^; “rttaCtrl” refers to Krt5-rtTA/+; En1^flox/flox^: “ko” refers to tetO-Cre/+;Krt5-rtTA/+; En1^flox/flox^. Alpha = 0.05.

**Table S4:**
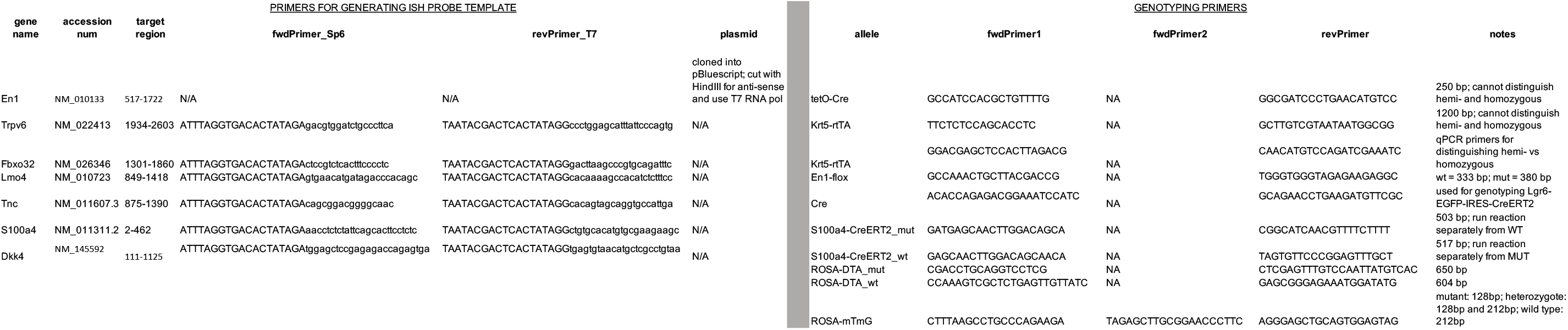
Sequences for polymerase chain reaction (PCR) primers used in this study.

## SUPPLEMENTAL DATA FILES

### Data S1

Tab delimited file of marker genes identified for all clusters from primary clustering

### Data S2

Tab delimited file of marker genes identified for all epidermal subclusters

### Data S3

Tab delimited file of marker genes identified for dermal subclusters

### Data S4

Tab delimited file containing all significant results of CellChat communication analysis

## REFERENCES

1. Kuno, Y. Human perspiration. (Thomas, 1956).

2. William Montagna & Paul F. Parakhal. The Structure and Function of Skin – 3rd Edition. (Elsevier Inc., 1974).

3. Kamberov, Y. G. et al. Comparative evidence for the independent evolution of hair and sweat gland traits in primates. Journal of Human Evolution 125, 99–105 (2018).

4. Lu, C. & Fuchs, E. Sweat Gland Progenitors in Development, Homeostasis, and Wound Repair. Cold Spring Harbor Perspectives in Medicine 4, a015222–a015222 (2014).

5. Cui, C.-Y. & Schlessinger, D. Eccrine sweat gland development and sweat secretion. Experimental dermatology 24, 644 (2015).

6. Montagna, W., Ellis, R. A. & Silver, A. Advances in Biology of Skin Vol. III, Eccrine Sweat Glands and Eccrine Sweating. (Pergamon Press Inc., 1962).

7. Bayuo, J. Management strategies of burns associated hyperthermia: A case report. Burns Open 1, 45–47 (2017).

8. Biggs, L. C. & Mikkola, M. L. Early inductive events in ectodermal appendage morphogenesis. Seminars in Cell & Developmental Biology 25–26, 11–21 (2014).

9. Lu, C. P. et al. Identification of Stem Cell Populations in Sweat Glands and Ducts: Roles in Homeostasis and Wound Repair. Cell 150, 136–150 (2012).

10. Cui, C.-Y., et al. Involvement of Wnt, Eda and Shh at defined stages of sweat gland development. Development 141, 3752–3760 (2014).

11. Taylor, D. K., Bubier, J. A., Silva, K. A. & Sundberg, J. P. Development, Structure, and Keratin Expression in C57BL/6J Mouse Eccrine Glands. Vet Pathol 49, 146– 154 (2012).

12. Lu, C. P., Polak, L., Keyes, B. E. & Fuchs, E. Spatiotemporal antagonism in mesenchymal-epithelial signaling in sweat versus hair fate decision. Science 354, aah6102 (2016).

13. Plikus, M. et al. Morpho-Regulation of Ectodermal Organs. The American Journal of Pathology 164, 1099–1114 (2004).

14. Andl, T., Reddy, S. T., Gaddapara, T. & Millar, S. E. WNT signals are required for the initiation of hair follicle development. Dev. Cell 2, 643–653 (2002).

15. Mayer, J. A., Foley, J., De La Cruz, D., Chuong, C.-M. & Widelitz, R. Conversion of the Nipple to Hair-Bearing Epithelia by Lowering Bone Morphogenetic Protein Pathway Activity at the Dermal-Epidermal Interface. The American Journal of Pathology 173, 1339–1348 (2008).

16. Kamberov, Y. G. et al. Modeling Recent Human Evolution in Mice by Expression of a Selected EDAR Variant. Cell 152, 691–702 (2013).

17. Headon, D. J. & Overbeek, P. A. Involvement of a novel Tnf receptor homologue in hair follicle induction. Nat. Genet. 22, 370–374 (1999).

18. Headon, D. J. et al. Gene defect in ectodermal dysplasia implicates a death domain adapter in development. Nature 414, 913–916 (2001).

19. Kamberov, Y. G. et al. A genetic basis of variation in eccrine sweat gland and hair follicle density. Proc. Natl. Acad. Sci. U.S.A. 112, 9932–9937 (2015).

20. Aldea, D. et al. The Transcription Factor Deaf1 Modulates Engrailed-1 Expression to Regulate Skin Appendage Fate. J Invest Dermatol 139, 2378–2381.e4 (2019).

21. Widelitz, R. B. & Chuong, C.-M. Early Events in Skin Appendage Formation: Induction of Epithelial Placodes and Condensation of Dermal Mesenchyme. Journal of Investigative Dermatology Symposium Proceedings 4, 302–306 (1999).

22. Huh, S.-H. et al. Fgf20 governs formation of primary and secondary dermal condensations in developing hair follicles. Genes Dev. 27, 450–458 (2013).

23. Biggs, L. C. et al. Hair follicle dermal condensation forms via Fgf20 primed cell cycle exit, cell motility, and aggregation. eLife 7, e36468 (2018).

24. Mok, K.-W. et al. Dermal Condensate Niche Fate Specification Occurs Prior to Formation and Is Placode Progenitor Dependent. Dev. Cell 48, 32–48.e5 (2019).

25. Gupta, K. et al. Single-Cell Analysis Reveals a Hair Follicle Dermal Niche Molecular Differentiation Trajectory that Begins Prior to Morphogenesis. Dev. Cell 48, 17–31.e6 (2019).

26. Loomis, C. A. et al. The mouse Engrailed-1 gene and ventral limb patterning. Nature 382, 360–363 (1996).

27. Aldea, D. et al. Differential modularity of the mammalian Engrailed 1 enhancer network directs sweat gland development. PLOS Genetics 19, e1010614 (2023).

28. Aldea, D., et al. Repeated mutation of a developmental enhancer contributed to human thermoregulatory evolution. PNAS 118, (2021).

29. Mainguy, G. et al. Regulation of Epidermal Bullous Pemphigoid Antigen 1 (BPAG1) Synthesis by Homeoprotein Transcription Factors. Journal of Investigative Dermatology 113, 643–650 (1999).

30. Glover, J. D. et al. The developmental basis of fingerprint pattern formation and variation. Cell 0, (2023).

31. Sgaier, S. K. et al. Genetic subdivision of the tectum and cerebellum into functionally related regions based on differential sensitivity to engrailed proteins. Development 134, 2325–2335 (2007).

32. Perl, A.-K. T., Wert, S. E., Nagy, A., Lobe, C. G. & Whitsett, J. A. Early restriction of peripheral and proximal cell lineages during formation of the lung. Proc. Natl. Acad. Sci. U.S.A. 99, 10482–10487 (2002).

33. Diamond, I., Owolabi, T., Marco, M., Lam, C. & Glick, A. Conditional gene expression in the epidermis of transgenic mice using the tetracycline-regulated transactivators tTA and rTA linked to the keratin 5 promoter. J. Invest. Dermatol. 115, 788–794 (2000).

34. Habib, N. et al. Massively parallel single-nucleus RNA-seq with DroNc-seq. Nat. Methods 14, 955–958 (2017).

35. Hu, P. et al. Dissecting Cell-Type Composition and Activity-Dependent Transcriptional State in Mammalian Brains by Massively Parallel Single-Nucleus RNA-Seq. Molecular Cell 68, 1006–1015.e7 (2017).

36. Grindberg, R. V. et al. RNA-sequencing from single nuclei. Proceedings of the National Academy of Sciences 110, 19802–19807 (2013).

37. Korsunsky, I. et al. Fast, sensitive and accurate integration of single-cell data with Harmony. Nat Methods 16, 1289–1296 (2019).

38. Hirai, Y., Nose, A., Kobayashi, S. & Takeichi, M. Expression and role of E- and P-cadherin adhesion molecules in embryonic histogenesis. I. Lung epithelial morphogenesis. Development 105, 263–270 (1989).

39. Mills, A. A. et al. p63 is a p53 homologue required for limb and epidermal morphogenesis. Nature 398, 708–713 (1999).

40. Pellegrini, G. et al. p63 identifies keratinocyte stem cells. Proceedings of the National Academy of Sciences 98, 3156–3161 (2001).

41. Driskell, R. R. et al. Distinct fibroblast lineages determine dermal architecture in skin development and repair. Nature 504, 277–281 (2013).

42. Bazzi, H., Fantauzzo, K. A., Richardson, G. D., Jahoda, C. A. B. & Christiano, A. M. The Wnt inhibitor, Dickkopf 4, is induced by canonical Wnt signaling during ectodermal appendage morphogenesis. Developmental Biology 305, 498–507 (2007).

43. Sick, S., Reinker, S., Timmer, J. & Schlake, T. WNT and DKK Determine Hair Follicle Spacing Through a Reaction-Diffusion Mechanism. Science 314, 1447–1450 (2006).

44. Fliniaux, I., Mikkola, M. L., Lefebvre, S. & Thesleff, I. Identification of dkk4 as a target of Eda-A1/Edar pathway reveals an unexpected role of ectodysplasin as inhibitor of Wnt signalling in ectodermal placodes. Developmental Biology 320, 60– 71 (2008).

45. Zhang, Y. et al. Reciprocal requirements for EDA/EDAR/NF-kappaB and Wnt/beta-catenin signaling pathways in hair follicle induction. Dev. Cell 17, 49–61 (2009).

46. Joyner, A. L. & Martin, G. R. En-1 and En-2, two mouse genes with sequence homology to the Drosophila engrailed gene: expression during embryogenesis. Genes Dev. 1, 29–38 (1987).

47. Jin, S. et al. Inference and analysis of cell-cell communication using CellChat. Nat Commun 12, 1088 (2021).

48. Voehringer, D., Liang, H.-E. & Locksley, R. M. Homeostasis and Effector Function of Lymphopenia-Induced “Memory-Like” T Cells in Constitutively T Cell-Depleted Mice1. The Journal of Immunology 180, 4742–4753 (2008).

49. Darwin, C. The variation of animals and plants under domestication. vol. II (John Murray, Albemarle Street, 1875).

50. Dhouailly, D. A new scenario for the evolutionary origin of hair, feather, and avian scales. J Anat 214, 587–606 (2009).

51. Zhang, Y. et al. Activation of beta-catenin signaling programs embryonic epidermis to hair follicle fate. Development 135, 2161–2172 (2008).

52. Mikkola, M. L. & Millar, S. E. The mammary bud as a skin appendage: unique and shared aspects of development. J Mammary Gland Biol Neoplasia 11, 187–203 (2006).

53. Chu, E. Y. et al. Canonical WNT signaling promotes mammary placode development and is essential for initiation of mammary gland morphogenesis. Development 131, 4819–4829 (2004).

54. Dhouailly, D. Dermo-epidermal interactions between birds and mammals: differentiation of cutaneous appendages. Development 30, 587–603 (1973).

55. Dhouailly, D. Regional specification of cutaneous appendages in mammals. Wilhelm Roux’ Archiv 181, 3–10 (1977).

56. Ferraris, C., Chevalier, G., Favier, B., Jahoda, C. A. & Dhouailly, D. Adult corneal epithelium basal cells possess the capacity to activate epidermal, pilosebaceous and sweat gland genetic programs in response to embryonic dermal stimuli. Development 127, 5487–5495 (2000).

57. Cygan, J. A., Johnson, R. L. & McMahon, A. P. Novel regulatory interactions revealed by studies of murine limb pattern in Wnt-7a and En-1 mutants. Development 124, 5021–5032 (1997).

58. GTEx Consortium. Human genomics. The Genotype-Tissue Expression (GTEx) pilot analysis: multitissue gene regulation in humans. Science 348, 648–660 (2015).

59. Reynolds, A. J. & Jahoda, C. A. Cultured dermal papilla cells induce follicle formation and hair growth by transdifferentiation of an adult epidermis. Development 115, 587–593 (1992).

60. Jahoda, C. A., Horne, K. A. & Oliver, R. F. Induction of hair growth by implantation of cultured dermal papilla cells. Nature 311, 560–562 (1984).

61. Edgar, R. Gene Expression Omnibus: NCBI gene expression and hybridization array data repository. Nucleic Acids Research 30, 207–210 (2002).

62. Snippert, H. J. et al. *Lgr6* Marks Stem Cells in the Hair Follicle That Generate All Cell Lineages of the Skin. Science 327, 1385–1389 (2010).

63. Muzumdar, M. D., Tasic, B., Miyamichi, K., Li, L. & Luo, L. A global double-fluorescent Cre reporter mouse. genesis 45, 593–605 (2007).

64. Schindelin, J. et al. Fiji: an open-source platform for biological-image analysis. Nat Methods 9, 676–682 (2012).

65. Macosko, E. Z., Basu, A., Satija, R., & others. Highly parallel genome-wide expression profiling of individual cells using nanoliter droplets. Cell 161, 1202–1214 (2015).

66. Hao, Y. et al. Integrated analysis of multimodal single-cell data. Cell 184, 3573–3587.e29 (2021).

67. Wu, T. et al. clusterProfiler 4.0: A universal enrichment tool for interpreting omics data. Innovation (Camb) 2, 100141 (2021).

68. Love, M. I., Huber, W. & Anders, S. Moderated estimation of fold change and dispersion for RNA-seq data with DESeq2. Genome Biol 15, 550 (2014).

69. Street, K. et al. Slingshot: cell lineage and pseudotime inference for single-cell transcriptomics. BMC Genomics 19, 477 (2018).

70. R Core Team. R: A Language and Environment for Statistical Computing. (R Foundation for Statistical Computing, 2022).

